# Ten3–Lphn2-mediated target selection across the extended hippocampal network demonstrates a repeated strategy for circuit assembly

**DOI:** 10.1101/2025.08.13.670207

**Authors:** Ellen C. Gingrich, Daniel T. Pederick, Yanbo Zhang, Liqun Luo

## Abstract

How do thousands of cell-surface proteins specify billions of neuronal connections in developing brains? We previously found that inverse expression of a ligand–receptor pair, teneurin-3 (Ten3) and latrophilin-2 (Lphn2) in CA1 and subiculum, instructs CA1→subiculum target selection through Ten3–Ten3 homophilic attraction and Ten3–Lphn2 heterophilic reciprocal repulsions. Here, we leveraged conditional knockouts to demonstrate that these mechanisms generalize to extended hippocampal networks, including entorhinal cortex and hypothalamus. Cooperation between attraction and repulsion differs depending on the order in which developing axons encounter the attractant and repellent subfields. Ten3 and Lphn2 can serve both as ligands for incoming axons and receptors for their own target selection, within the same neuron; Ten3 can be repulsive or attractive as ligand or receptor. Thus, multifunctionality and repeated use, together with recurrent circuit motifs prevalent in the brain, enable one ligand–receptor pair to instruct target selection of many more neurons.

## INTRODUCTION

During development, neural circuits self-assemble with exquisite wiring precision to enable proper functions such as memory and spatial navigation. Roger Sperry proposed in his chemoaffinity hypothesis that, to accomplish this feat, individual neurons carry unique molecular identification tags that distinguish themselves from neighboring neurons.^1^ Many cell-surface proteins (CSPs) have since been identified as critical regulators of axon guidance and target selection.^2,3^ These CSPs can be expressed on both pre- and post-synaptic cells and use attraction and repulsion to specify neuronal partner matching.^4–13^

The ∼10^11^ neurons and 10^14^ synaptic connections in the human brain vastly outnumber the 3000 CSPs available to instruct them,^14–17^. Among multiple strategies that have been proposed to overcome this discrepancy, one is to re-use the same CSPs at spatially distinct locations.^18^ For example, the ligand–receptor pair ephrin-A and EphA display inverse expression in several retinotopic nodes of the visual circuit including the retina, superior colliculus (SC), lateral geniculate nucleus (LGN), and the primary visual cortex (V1).^19^ In the retina→SC connection, ephrin-A–EphA-mediated reciprocal repulsion during target selection has been shown to establish the precise anterior–posterior retinotopy.^5,6,20–22^ However, although ephrin-A–EphA signaling is hypothesized to directly regulate retinotopic connectivity across other nodes, this has not been studied extensively beyond retina→SC and the use of whole animal knockouts cannot distinguish their roles in axons vs targets.^23–25^ The only study that employed conditional knockout (of ephrin-A5) we are aware of uncovered the importance of target-independent axon-axon interactions in establishing retinocollicular connectivity.^26^

Compared to sensory and motor systems, much less is known about wiring mechanisms of central brain circuits. One such circuit with well-characterized topographic projections across multiple nodes is the extended hippocampal network (**Figure 1A_1–2_**). The medial and lateral hippocampal subnetworks (MHN and LHN) preferentially process spatial and object-related information, respectively.^27,28^ The MHN consists, in part, of interconnected medial entorhinal cortex (MEC), proximal CA1 (pCA1), distal subiculum (dSub), and lateral subdivision of the medial mammillary nucleus (lat-mMN) of the hypothalamus. Conversely, the LHN consists of interconnected lateral entorhinal cortex (LEC), distal CA1 (dCA1), proximal subiculum (pSub), and medial mMN (med-mMN) (**Figure 1A**).^29–35^

**Figure 1.**
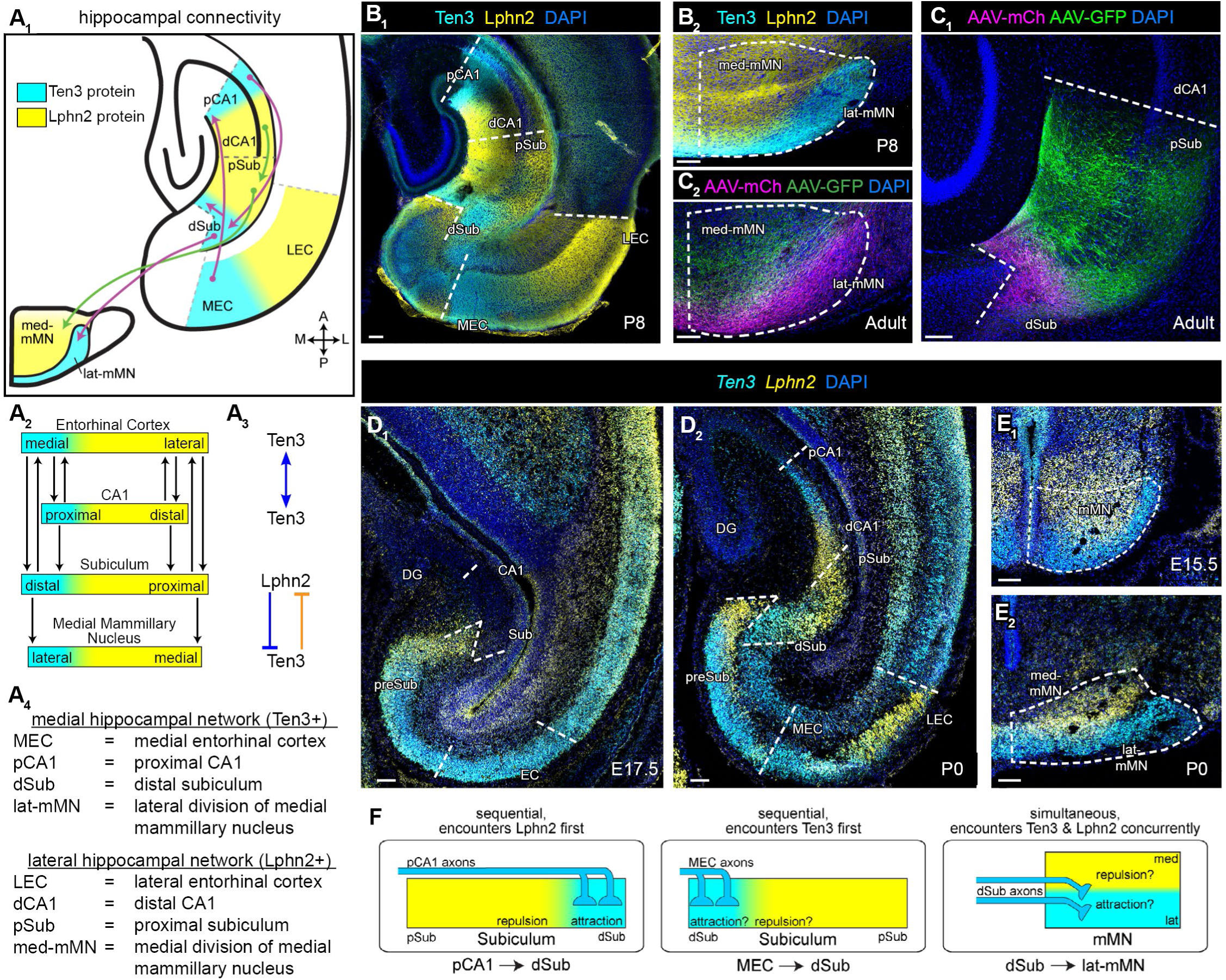
Inverse Ten3 and Lphn2 expression patterns match extended hippocampal network connectivity. (A) Overview. (A_1_) Summary of topographic connections studied, overlaid on P8 Ten3 and Lphn2 protein expression. (A_2_) All connectivity between four nodes of the medial (cyan) and lateral (yellow) hippocampal network. (A_3_) Summary of molecular interactions between Ten3 and Lphn2. The bidirectional arrow represents attraction; **––|** represents repulsion from target to axon. Blue mechanisms are used for Ten3^+^ axons and orange mechanism is used for Lphn2^+^ axons. (A_4_) Anatomical abbreviations. (B) Inverse protein expression of Ten3 and Lphn2 in hippocampus and entorhinal cortex (B_1_) and medial mammillary nucleus (mMN) of the hypothalamus (B_2_) of P8 *Ten3^HA/HA^;Lphn2^mVenus/mVenus^*mouse. (C) Dual anterograde tracer injections into neighboring sites in distal subiculum (dSub) and proximal subiculum (pSub; C_1_) result in topographic projections to neighboring sites in lat-mMN and med-mMN, respectively (C_2_). (D) mRNA expression of *Ten3* (cyan) and *Lphn2* (yellow) at E17.5 (D_1_) and P0 (D_2_) in hippocampus and entorhinal cortex. At E17.5, only *Ten3* mRNA is detectable in EC. By P0, CA1, Sub, and EC all have inverse mRNA expression of *Ten3* and *Lphn2*. (E) mRNA expression of *Ten3* (cyan) and *Lphn2* (yellow) at E15.5 (E_1_) and P0 (E_2_) in mMN. (F) Summary of three different axon entry routes with respect to Ten3 and Lphn2 expression subfields in target. Scale bars, 100 μm. See Figure S1 for additional data. In this and all subsequent figures, A, anterior; P, posterior; M, medial; L, lateral.

We previously found a CSP pair, teneurin-3 (Ten3) and latrophilin-2 (Lphn2), that exhibit inverse expression in both CA1 and subiculum, such that the interconnected regions of pCA1→dSub and dCA1→pSub express Ten3 and Lphn2, respectively (**Figure 1A_1-2_**).^8,9^ Loss- and gain-of-function analyses revealed that the CA1→subiculum specificity is produced by the cooperation between homophilic Ten3–Ten3-mediated attraction and heterophilic Ten3–Lphn2-mediated reciprocal repulsions (**Figure 1A_3_**).^9,36^ Given that Ten3 and Lphn2 also exhibit inverse expression in other nodes of the MHN and LHN including entorhinal cortex (EC) and medial mammillary nucleus (mMN),^8,9,37,38^ here we address whether other parts of the network follow a ‘Ten3→Ten3, Lphn2→Lphn2’ connectivity rule and, if so, whether this ligand–receptor pair is used repeatedly to instruct the precise assembly at other nodes of the network using systematic conditional knockout analyses.

## RESULTS

### The extended hippocampal network follows a ‘Ten3→Ten3, Lphn2→Lphn2’ connectivity rule

Beyond its inverse expression in CA1 and subiculum, Ten3 is also enriched in subfields of several other nodes in the extended hippocampal network, including MEC and lat-mMN.^8,38^ Conversely, Lphn2 is enriched in LEC (**Figure 1A**).^9,37,38^ Immunostaining for tagged Ten3 and Lphn2 (in *Ten3^HA/HA^;Lphn2^mVenus/mVenus^* knockin mice^39,40^) confirmed the striking inverse expression of these two proteins at postnatal day 8 (P8; **Figure 1B**). As previously reported, Ten3 protein is enriched in pCA1, dSub, MEC, and lat-mMN; by contrast, Lphn2 protein is enriched in dCA1, pSub, LEC, and med-mMN (**Figure 1A, B**).^8,9,37^ Given this topographically-restricted expression, we considered whether these two CSPs may serve a broad role in assembling the extended hippocampal networks. Previous studies have focused on the relationship between Ten3–Lphn2 inverse expression and connection topography in the hippocampal-entorhinal circuitry.^8,9,37^ To investigate this relationship in hippocampal outputs to the hypothalamus, we injected adeno-associated viruses (AAVs) as anterograde tracers into neighboring locations in the subiculum (*AAV-GFP* into Lphn2-high pSub, *AAV-mCherry* (*mCh*) into Ten3-high dSub) and assessed their projections to mMN (**Figure 1C_1_**, **Figure S1A_1_**). Note that in this study, pSub refers to the entirety of the Lphn2-high region of subiculum which anatomically constitutes approximately 75% of the length of subiculum (**Figure S1A_1_**).^9^ We found that GFP^+^ axons were restricted to Lphn2-high med-mMN whereas mCh^+^ axons filled Ten3-high lat-mMN (**Figure 1C_2_**, **Figure S1A_2,_ B**), validating that the ‘Ten3→Ten3, Lphn2→Lphn2’ connectivity rule also applies to subiculum→mMN projection, analogous to projections between CA1, subiculum, and EC (**Figure 1A_1, 2_**).^8,9^ Of note, the axons from subiculum to mMN did not appear segregated, instead forming a uniform bundle that splits into the appropriate subfield as it enters the mMN target region (**Figure S1B**).

### Spatiotemporal patterns of Ten3 and Lphn2 expression with respect to target entry of developing axons

To instruct target selection of axons connecting these nodes, Ten3 and Lphn2 must be inversely expressed at each node prior to the maturation of the circuit. To establish expression onset, we performed dual *in situ* hybridization for *Ten3* and *Lphn2* mRNA using RNAScope^41^ across multiple developmental timepoints. At E15.5, *Ten3* and *Lphn2* were already inversely expressed in mMN (**Figure 1E**) but exhibited minimal expression in the hippocampal and entorhinal regions. By E17.5, *Ten3* was highly expressed in entorhinal cortex whereas *Lphn2* expression remained low (**Figure 1D_1_**), marking this as the only case we observed where *Ten3–Lphn2* were not inversely expressed at onset. By P0, CA1, subiculum, and entorhinal cortex all exhibited inverse expression, resembling protein staining at P8 (**Figure 1B_1_, D_2_**) and in previous studies.^9,38^ *Ex vivo* tracing from MEC showed that axons began to invade dSub at E17.5 and increased in density through P0 (**Figure S1C, D**), roughly aligning with the onset of *Ten3* and *Lphn2* expression in dSub and pSub, respectively. Although little is known about the timing of subiculum→mMN axon targeting, expression of *Ten3* and *Lphn2* in the subiculum axon bundle in the whole-brain staining described in our companion manuscript^39^ indicated that these axons reach mMN around P0. Since each node of the circuit expressed *Ten3* and *Lphn2* mRNA (**Figure 1D, E**), the protein distribution we observed (**Figure 1B**) is likely contributed to by both input axons and target neurons, further reinforcing the prevalence of like–like connectivity.

Although each of the four nodes we examined—CA1, subiculum, entorhinal cortex, and mMN—exhibited striking inverse expression of Ten3 and Lphn2, the spatial organization of developing axons encountering each target node in relation to the expression pattern could influence target selection strategy. By re-examining our previous study,^8^ characterizing adult axon entry routes,^42–44^ performing developmental tracing, and cataloguing Ten3–Lphn2 protein expression, we established three distinct spatial configurations in which axons may encounter Ten3 and Lphn2 subfields in the target (**Figure 1F**; **Figure S1B, E–G**). (1) For CA1→subiculum at P2, Ten3^+^ pCA1 axons first encounter the Lphn2^+^ pSub, then the Ten3^+^ dSub (**Figure 1F, left**)^8^, consistent with adult single axon tracing studies.^43,44^ (2) For entorhinal cortex→subiculum, Ten3^+^ MEC axons first encountered the Ten3^+^ dSub, then the Lphn2^+^ pSub, as apparent in *ex vivo* tracing of MEC in P0 *Ten3^HA/HA^;Lphn2^mV/mV^* pups (**Figure 1F, middle**; **Figure S1E**). This is also consistent with adult single axon tracing from entorhinal cortex.^42,45,46^ (3) For subiculum→mMN, Ten3^+^ dSub axons encountered the Ten3^+^ lat-mMN and Lphn2^+^ med-mMN subfields simultaneously, confronting them with a binary choice (**Figure 1F, right**; **Figure S1B, F, G**). Likewise, Lphn2^+^ pSub axons encountered Ten3^+^ and Lphn2^+^ subfields simultaneously (**Figure S1B, F, G**). Our previous work has established that both Ten3–Ten3 homophilic attraction and Ten3–Lphn2 heterophilic reciprocal repulsion are critical in the first scenario (**Figure 1F, left**). However, it remains unclear how the other two configurations of developing axons encountering the Ten3^+^ or Lphn2^+^ target subfields (**Figure 1F, middle and right**) may influence the use of these mechanisms.

### Ten3 is required in MEC axons for their precise targeting to dSub

Ten3–Ten3 homophilic attraction and Lphn2–Ten3 heterophilic repulsion are critical for establishing the appropriate topography of the pCA1→dSub connection (**Figure 1A_3_**).^8,9^ To determine if these mechanisms are re-used at the MEC→dSub connection, we first asked if Ten3 is required in MEC axons for targeting using a conditional knockout strategy. We injected *Cre-GFP*-expressing lentivirus into MEC of postnatal day 0 (P0) wildtype or *Ten3^fl/fl^* pups, titrating the concentration to allow for a roughly even proportion of Cre^+^ and Cre^–^neurons (**Figure 2A, left**). At P42 or older, we injected a mix of anterograde *AAV-CreON-GFP* and *AAV-CreOFF-mCh* into MEC of these mice (**Figure 2A, middle**) to assess the targeting of control and *Ten3*-null axons within the same animal (**Figure 2A, right**). Injections were restricted to within the most medial 20% of entorhinal cortex to ensure tracing of mostly Ten3^+^ axons (**Figure 2B**). See **Table S1** for detailed information on genotypes and injection conditions.

**Figure 2.**
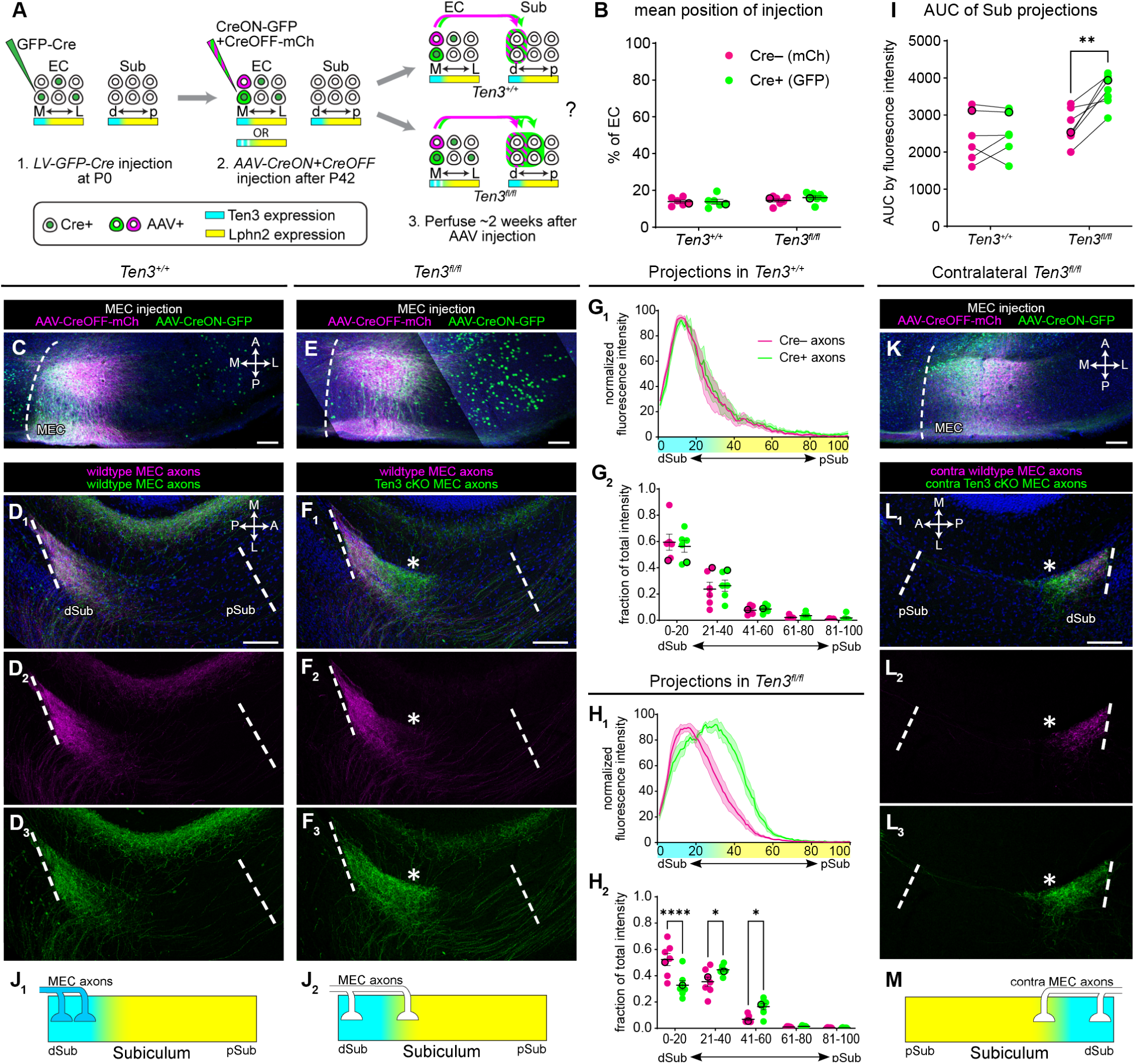
Ten3 is required in MEC axons for their precise targeting to distal subiculum. (A) Injection strategy and possible results for tracing MEC axons into subiculum of control (top right) and *Ten3^fl/fl^* mice (bottom right; white lines in cyan area in the bottom designate mosaic *Ten3* deletion in MEC neurons). (B) Mean injection site positions along the MEC-to-LEC axis show no differences between *Ten3^+/+^* controls (n = 6) and *Ten3^fl/fl^* (n = 7) mice. Black outline indicates representative animals shown in C, E. Mean ± SEM. Mann-Whitney tests corrected for multiple comparisons (Holm-Šídák). (C) Representative image of the *AAV-CreON/CreOFF* injection site of *Ten3^+/+^* controls show both viruses restricted to the most medial region of entorhinal cortex. Same animal as in D. (D) Representative images of projection of Cre^–^ axons (magenta) and Cre^+^ axons (green) into molecular layer of subiculum in *Ten3^+/+^* controls. The three images are from the same confocal section showing merge (D_1_), magenta only (D_2_), or green only (D_3_) channels. (E, F) Same as C, D but for *Ten3^fl/fl^* mice. *Ten3^MEC^*-cKO axons spread beyond the Cre^–^ control axons toward pSub (asterisk in F, in identical position across three images). Green dots in C and E come from *Cre-GFP* lentivirus. (G_1_) Normalized fluorescence intensity traces of the Cre^–^ (magenta) and Cre^+^ (green) projections along the dSub–pSub axis in *Ten3^+/+^* controls (n = 6 mice). Ten3 (cyan) and Lphn2 (yellow) expression data is represented along the x-axis. Mean (dark line) ± SEM (shaded area). (G_2_) Fraction of total projection intensity (same data as G_1_) in 20% bins of subiculum axis of the Cre^–^ (magenta) and Cre^+^ (green) projections along the dSub–pSub axis in *Ten3^+/+^* controls (n = 6 mice). Black outlines indicate representative animal shown in D. Mean ± SEM. Two-way ANOVA corrected for multiple comparisons (Šídák correction), no significant differences. (H) Same as G but for *Ten3^fl/fl^* (n = 7) mice. **** p < 0.0001, * p < 0.05. (I) Area under the curve of averaged subiculum projections for *Ten3^+/+^* controls (n = 6) and *Ten3^fl/fl^* (n = 7) mice. Lines are between Cre^–^ (magenta) and Cre^+^ (green) projections within the same animal. Black outlines indicate representative animals shown in D, F. Mann-Whitney tests corrected for multiple comparisons (Holm-Šídák), ** p < 0.01. (J) Summary of target selection of MEC axons in subiculum in control (J_1_) and *Ten3^MEC^*-cKO (J_2_) based on results in C–I. (K, L) Same as E, F, but for projections into contralateral subiculum. Cre^+^, *Ten3^MEC^*-cKO axons spread beyond the Cre^–^ control axons toward pSub (asterisk, in identical position across three images). (M) Summary of target selection of *Ten3^MEC^*-cKO axons in contralateral subiculum. Scale bars, 100 μm. See Figure S2 for additional data.

In *Ten3^+/+^* controls, mCh^+^ (Cre^–^) and GFP^+^ (Cre^+^) MEC neurons alike exclusively projected to the distal, Ten3-high region of the subiculum (**Figure 2C, D, G**). In *Ten3^fl/fl^* mice, mCh^+^ axons (Cre^–^, functionally wildtype) remained restricted to Ten3-high dSub; however, GFP^+^ (Cre^+^, *Ten3* conditionally knocked out, or *Ten3^MEC^-cKO*) axons invaded Lphn2-high regions of pSub (**Figure 2E, F, H**). Quantification of axon fluorescence intensity (FI) showed that only GFP^+^ axons in *Ten3^fl/fl^* mice had a proximal shift along the distal–proximal axis of the subiculum (**Figure 2G_1_, H_1_**). This shift resulted in lower axon density in the appropriate Ten3-high dSub and increased density in the Lphn2-high pSub (**Figure 2G_2_, H_2_**). This mistargeting also caused GFP^+^ axons to cover a larger region of subiculum, as evidenced by area under the curve of FI traces, when Cre^–^ and Cre^+^ projections were compared pairwise within animals (**Figure 2I**). A similar phenotype was observed for target selection of MEC axons in CA1: GFP^+^ (and *Ten3^MEC^-cKO*) axons in *Ten3^fl/fl^* mice spread more distally into Lphn2-high regions compared to mCh^+^ wildtype axons (**Figure S2A–D**).

MEC axons also have topographic projections to contralateral dSub and pCA1 that could be affected by *Ten3^MEC^-cKO*. Although these projections are much sparser than the ipsilateral connection and were not observable in every animal, GFP^+^ axons also mistargeted into Lphn2-high regions of pSub and dCA1 in *Ten3^fl/fl^* mice compared to internal control mCh^+^ axons (**Figure 2K, L**; **Figure S2F, H**). These observations suggest that ipsi- and contralateral projections use similar mechanisms for subfield selection. Taken together, these data demonstrated that Ten3 is required in MEC neurons for axons to precisely target to dSub and pCA1 (**Figure 2J, M**; **Figure S2E, G**).

### Ten3 and Lphn2 in subiculum cooperate to instruct the precise targeting of MEC axons

We next asked whether Ten3 and/or Lphn2 are required in the subiculum target neurons, in addition to requiring Ten3 in MEC axons, for the MEC→dSub projection. We used a pan-subiculum Cre driver, *Nts-Cre*, which exhibits dense expression in subiculum at P0 (**Figure 3A**; **Figure S3A**) but minimal expression in entorhinal cortex or CA1 (**Figure S3A, B**), to conditionally knockout *Ten3*, *Lphn2*, or both selectively in subiculum.

**Figure 3.**
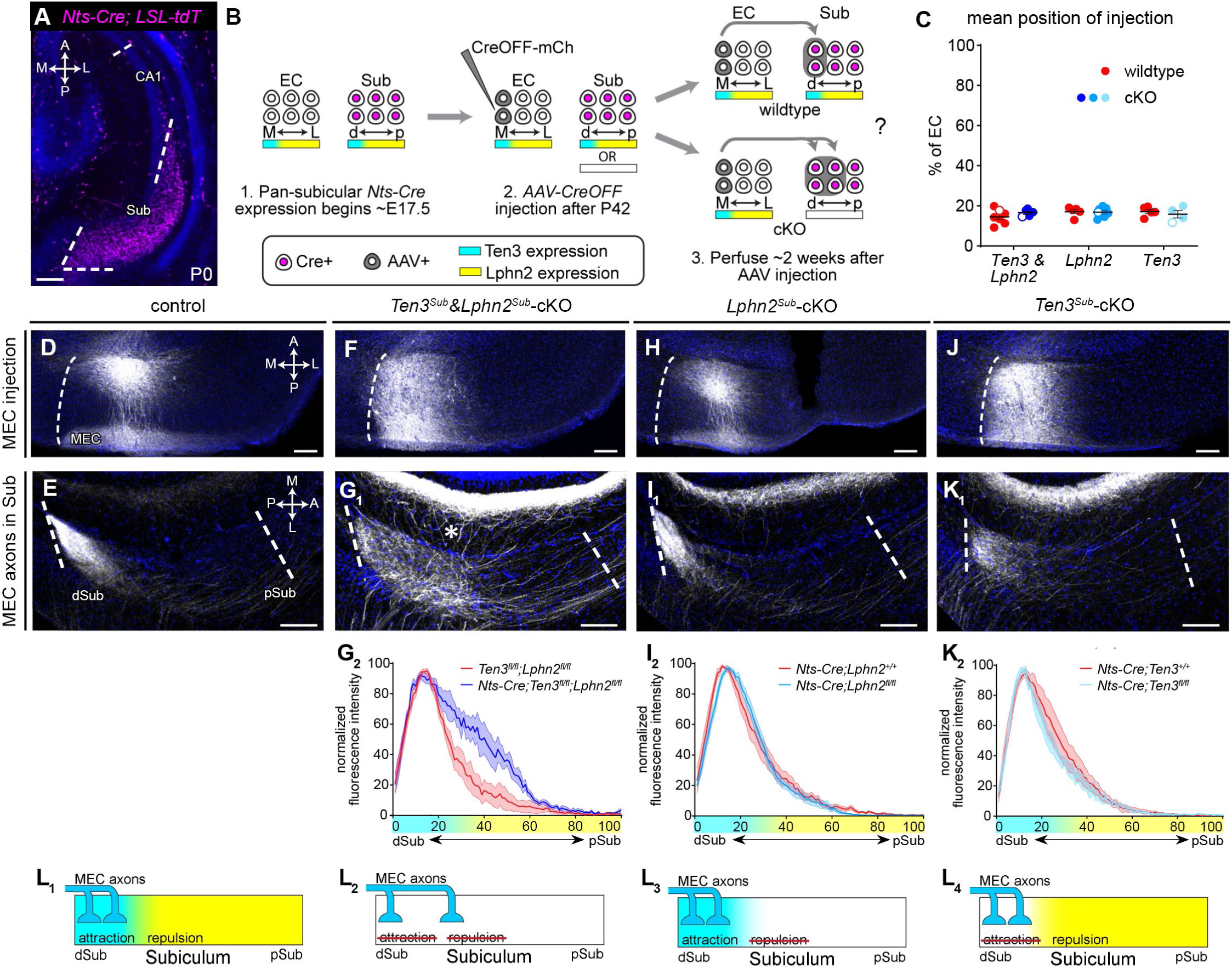
Ten3 and Lphn2 in subiculum cooperate to instruct the precise targeting of MEC axons. (A) *Nts-Cre* is densely expressed in subiculum at P0, assessed by a nuclear tdTomato reporter. Scale bar, 150 μm. (B) Injection strategy and possible results for tracing MEC axons into subiculum of control (top right) and *Ten3^Sub^&Lphn2^Sub^-*cKO mice (bottom right). (C) Mean positions of injection sites along the MEC-to-LEC axis show no differences between *Ten3^Sub^&Lphn2^Sub^-*cKO*, Lphn2^Sub^-*cKO, *Ten3^Sub^*-cKO (n = 5, 7, 4 mice, respectively), and their respective littermate controls (n = 7, 5, 5). Open circles indicate representative animals shown in D, F, H, J. Mean ± SEM. Mann-Whitney tests corrected for multiple comparisons (Holm-Šídák). (D) Representative image of the *AAV-CreOFF* injection site of *Ten3^fl/fl^;Lphn2^fl/fl^* controls shows that virus is restricted to the most medial region of entorhinal cortex. Same animal as in E. Scale bar, 100 μm. (E) Representative image of projection into molecular layer of subiculum for *Ten3^fl/fl^;Lphn2^fl/fl^* (control). Scale bar, 100 μm. (F, H, J) Same as D, but for genotypes indicated above. (G_1_, I_1_, K_1_) Same as E, but for genotypes indicated above. MEC axons in *Ten3^Sub^&Lphn2^Sub^*-cKO mice spread proximally beyond dSub (asterisk in G_1_). (G_2_, I_2_, K_2_) Normalized fluorescence intensity traces in genotypes as indicated. Mean (dark line) ± SEM (shaded area). n = 7, 5 for control and *Ten3^Sub^&Lphn2^Sub^-*cKO mice, respectively (G_2_); n = 5, 7 for control and *Lphn2^Sub^*-cKO mice, respectively (I_2_); n = 5, 4 for control and *Ten3^Sub^*-cKO mice, respectively (K_2_). Ten3 (cyan) and Lphn2 (yellow) expression data is represented along the x-axis. Mean (dark line) ± SEM (shaded area). (L) Summary of target selection of MEC axons in subiculum in control (L_1_) or when *Ten3* (L_4_), *Lphn2* (L_3_), or both (L_2_) were conditionally knocked out in subiculum target. See Figures S3–S4 for additional data.

We labeled MEC→dSub axon projections by injecting *AAV-CreOFF-mCh* into MEC (**Figure 3B, C**). To examine if target deletions could recapitulate the phenotype observed in *Ten3^MEC^-cKO* (**Figure 2J**; which presumably lose both Ten3–Ten3 attraction and Lphn2–Ten3 repulsion), we started by assessing axon distributions in *Ten3* and *Lphn2* double cKO mice. In *Nts-Cre;Ten3^fl/fl^;Lphn2^fl/fl^* (*Ten3^Sub^&Lphn2^Sub^-*cKO) mice, we observed mistargeting of Ten3^+^ axons into the Lphn2-high subfield of subiculum compared to *Ten3^fl/fl^;Lphn2^fl/fl^*littermate controls (**Figure 3D–G**). By contrast, we did not observe mistargeting of MEC→pCA1 axon in the same experiment (**Figure S4A–C**), reinforcing the specificity of *Nts-Cre* for subiculum-restricted manipulations. These experiments demonstrated that Ten3 and/or Lphn2 are required in the subiculum target for Ten3^+^ MEC axons to target to Ten3^+^ dSub, through Ten3–Ten3 homophilic attraction, Lphn2–Ten3 heterophilic repulsion, or a combination.

To examine the relative contribution of Ten3–Ten3 homophilic attraction and Lphn2–Ten3 heterophilic repulsion, we repeated these experiments with single cKOs: (1) *Nts-Cre;Lphn2^fl/fl^* (*Lphn2^Sub^-*cKO; with *Nts-Cre;Lphn2^+/+^* littermate controls); and (2) *Nts-Cre;Ten3^fl/fl^* (*Ten3^Sub^-*cKO; with *Nts-Cre;Ten3^+/+^* littermate controls). As expected, CA1 targeting was normal under both conditions (**Figure S4D, E**). Unexpectedly, however, Ten3^+^ MEC axons appeared to target the appropriate subiculum region in both single knockout conditions (**Figure 3H–K**).

These data suggest that Ten3–Ten3-mediated attraction and Lphn2–Ten3-mediated repulsion are largely redundant, with each mechanism compensating for the loss of the other (**Figure 3L**). This is contrary to our previous studies of pCA1→dSub targeting, where conditional knockout of either *Ten3* (attraction) or *Lphn2* (repulsion) in the subiculum target produces mistargeting phenotypes.^8,9^ One possibility is that, unlike the pCA1→dSub projection where axons encounter a large Lphn2-high region of pSub before contacting the appropriate dSub target, in the MEC→dSub projection, MEC axons projecting to dSub encounter the Ten3 attractant first (**Figure 1F**, **Figure S1E**), such that attraction alone may stabilize MEC axons before they reach the Lphn2-high pSub. Conversely, even without attraction, the repulsive forces emanating from Lphn2 are sufficient to preclude axons from entering pSub. Only when both Ten3 and Lphn2 are removed from the target do MEC axons phenocopy loss of Ten3 in axons, supporting that Ten3–Ten3 attraction and Lphn2–Ten3 repulsion cooperate in mediating MEC→dSub target selection.

### Ten3 is required in subiculum for dSub axons to precisely target to lateral mMN

Our data reveal that the precise targeting of pCA1→dSub and MEC→dSub both require the cooperation of Ten3–Ten3 attraction and Lphn2–Ten3 repulsion, but differ in their degree of reliance on these mechanisms individually, in accordance with the order in which Ten3^+^ axons encounter Ten3^+^ or Lphn2^+^ target subfields. We next asked what happens during a third scenario where axons encounter Ten3 and Lphn2 concurrently, as seen in dSub→lat-mMN connection (**Figure 1F**; **Figure S1F, G**).

We first assessed if Ten3 is required in dSub axons for appropriate targeting to lat-mMN by tracing dSub axons in *Ten3^Sub^*-cKO mice using the same *Nts-Cre* driver (low mMN expression; **Figure S3C**) as above (**Figure 4A, B**). Note that unlike in **Figure 3**, where *Ten3^Sub^*-cKO was used to examine the consequence of <u>target</u> deletion for MEC→dSub projection, hereafter we leveraged *Ten3^Sub^*-cKO to examine the consequence of <u>axon</u> deletion for subiculum→mMN projection. In control animals, dSub axons were restricted to the Ten3-high lateral region of mMN (lat-mMN; **Figure 4C, D**). However, in *Ten3^Sub^*-cKO animals, some dSub axons spread into Lphn2-high med-mMN (**Figure 4E, F**). We quantified axon mistargeting by defining the lat-mMN and med-mMN boundaries using DAPI counterstaining **(Figure S5H)** and calculating the fraction of fluorescence intensity in med-mMN over total fluorescence intensity. In control animals, axons were predominantly restricted to the lat-mMN, whereas in *Ten3^Sub^*-cKO, the fraction of axons in med-mMN significantly increased (**Figure 4G**). Mistargeting of dSub axons in *Ten3^Sub^*-cKO mice were also observed in an independent cohort of animals using an alternative analysis method **(Figure S5A–F)**. This sparse labeling of *Ten3*-null axons showed that individual axons did not exclusively mistarget to just the neighboring Lphn2-high subfield, unlike in MEC→dSub. We also observed this dSub axonal spread to med-mMN as these projections cross the midline into contralateral mMN, suggesting that once the targeting is disrupted, the axons remain in the incorrect subfield (**Figure 4I; Figure S5F_2,_ I**). Taken together, these data demonstrate that Ten3 is required in dSub axons to precisely target lat-mMN (**Figure 4H, S5G**).

**Figure 4.**
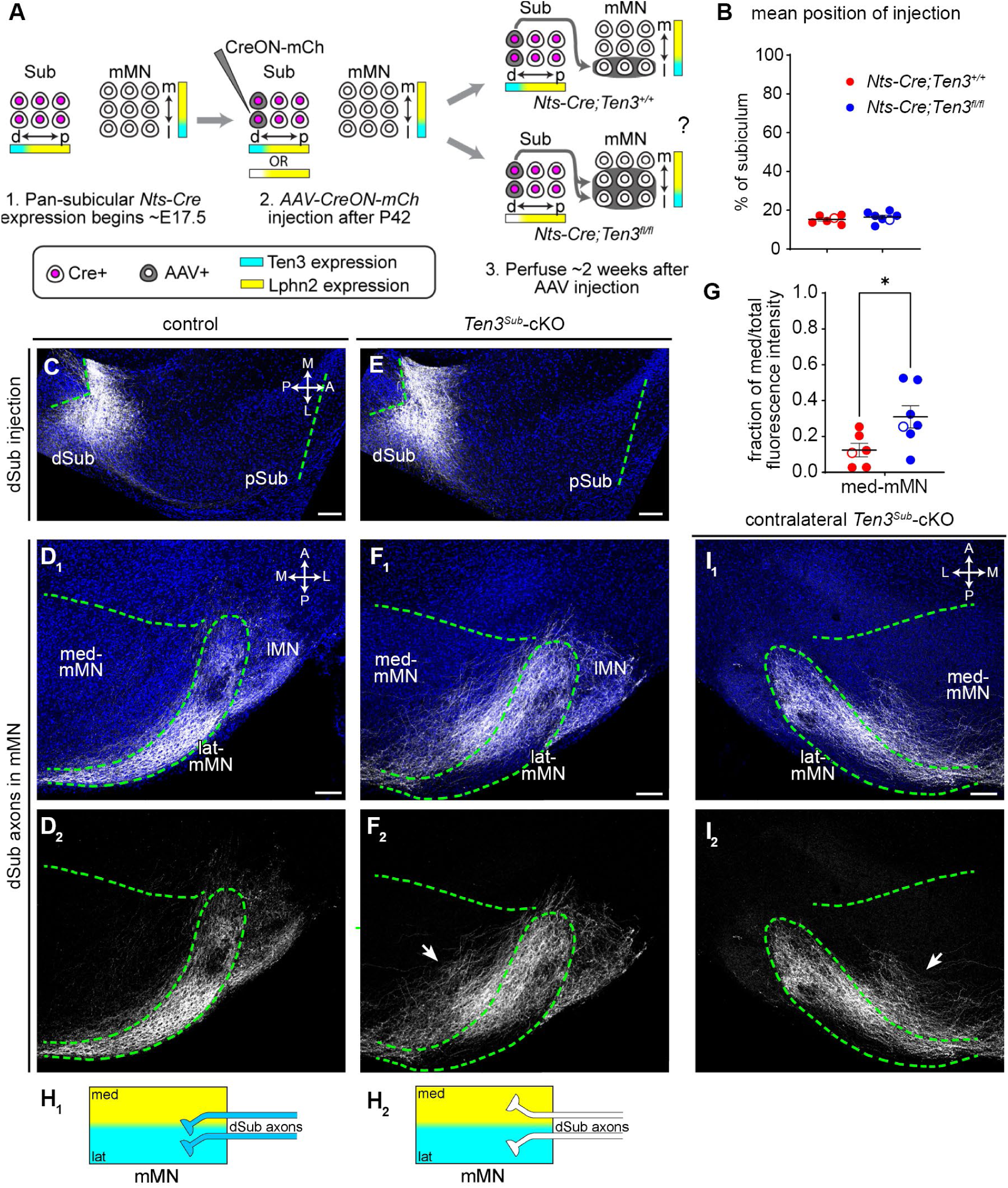
Ten3 is required in subiculum for dSub axons to precisely targeting to lateral mMN. (A) Injection strategy and possible results for tracing dSub axons into mMN of control (top right) and *Ten3^Sub^*-cKO mice (bottom right). (B) Mean positions of injection sites along the dSub-to-pSub axis show no differences between controls (n = 6) and *Ten3^Sub^*-cKO (n = 7) mice. Open circles indicate representative animals shown in C, E. Mean ± SEM. Mann-Whitney test. (C) Representative image of the *AAV-mCh* (gray) injection site of controls show that the virus is restricted to the most distal region of subiculum. Same animal as in D. (D) Representative images of projection of dSub axons (gray) into mMN of controls (D_1_). Bottom panel shows axons without DAPI counterstain (D_2_). Labeling in the lateral mammillary nucleus (lMN) are unrelated axons from pre-subiculum. (E, F) Same as C, D, but for *Ten3^Sub^*-cKO mice. *Ten3^Sub^*-cKO axons spread into the med-mMN (arrow). (G) Fraction of total projection intensity of the dSub axons in med-mMN in controls (red, n = 6) and *Ten3^Sub^*-cKO (blue, n = 7). Open circles indicate representative animals shown in D, F. Mean ± SEM. Mann-Whitney test, * p < 0.05. (H) Schematic summary of target selection of dSub axons in mMN of control (H_1_) and *Ten3^Sub^*-cKO (H_2_) mice based on results in C–G. (I) Same as F, but for projections into contralateral mMN. Contralateral *Ten3^Sub^*-cKO axons spread into the med-mMN (arrow). Scale bars, 100 μm. See Figures S3, S5, and S7 for additional data.

### Lphn2 is required in mMN for the precise targeting of dSub axons to lat-mMN

Given that Ten3 is required in dSub axons for them to appropriately target to mMN, we next asked whether Ten3 and/or Lphn2 are also required in the mMN target. To do this, we used *Sim1-Cre*, a Cre driver with dense expression in mMN as early as E15.5 (**Figure 5A**, **Figure S3D**), but minimal expression in subiculum (**Figure S3E**).

**Figure 5.**
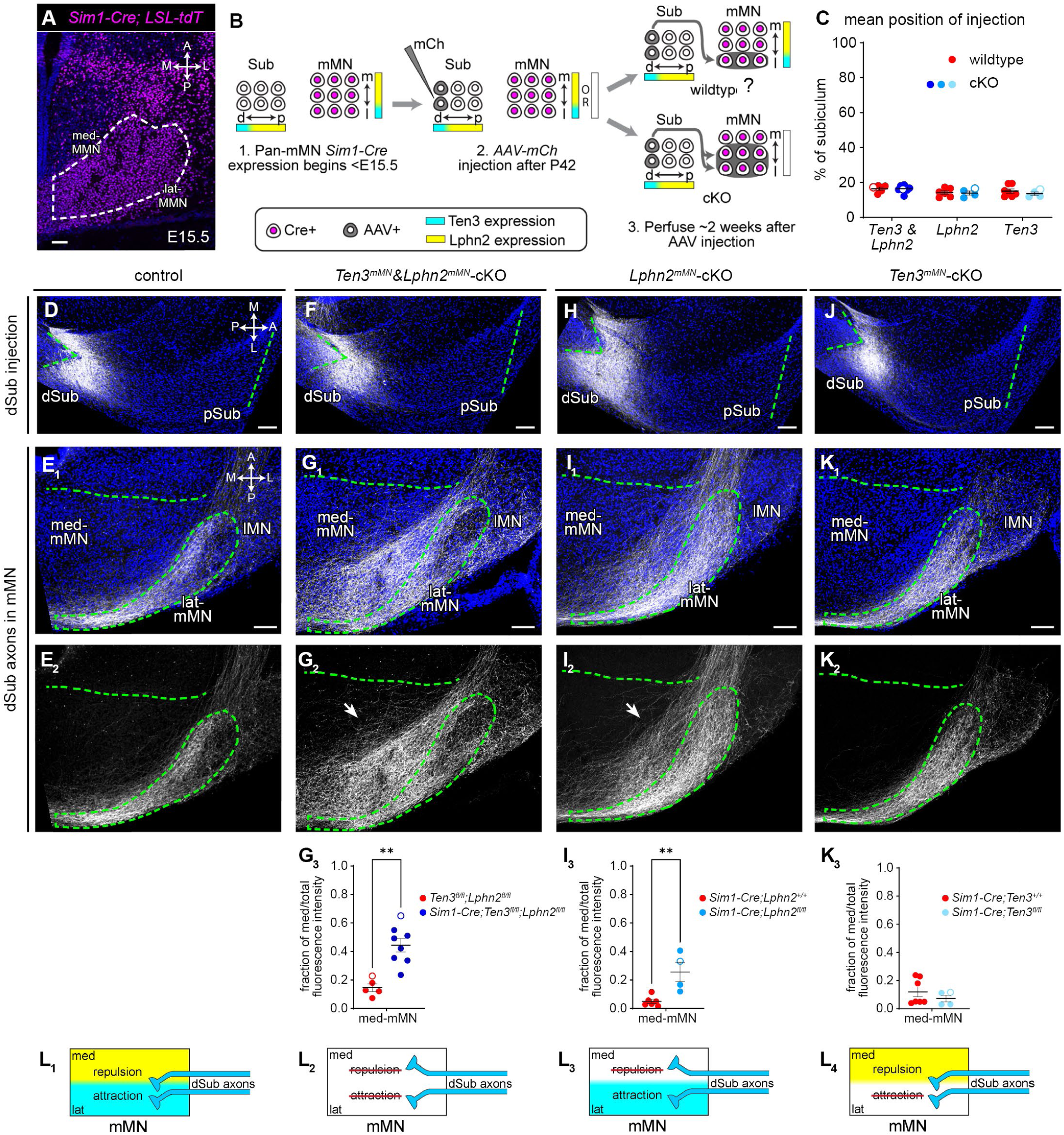
Lphn2 is required in mMN for the precise targeting of dSub axons to lat-mMN. (A) *Sim1-Cre* is densely expressed in mMN by E15.5, assessed by a nuclear tdTomato reporter. Scale bar, 50 μm. (B) Injection strategy and possible results for tracing dSub axons into mMN of controls (top right) and *Ten3^mMN^&Lphn2^mMN^-*cKO mice (bottom right). (C) Mean positions of injection sites along the dSub-to-pSub axis show no significant differences between *Ten3^mMN^&Lphn2^mMN^-*cKO, *Lphn2^mMN^-*cKO, *and Ten3^mMN^-*cKO (n = 8, 4, 4 mice respectively), and their respective littermate controls (n = 5, 6, 7, respectively). Open circles indicate representative animals shown in D, F, H, J. Mean ± SEM. Mann-Whitney tests corrected for multiple comparisons (Holm-Šídák). (D) Representative image of the *AAV-mCh* (gray) injection site of controls show that the virus is restricted to the most distal region of subiculum. Same animal as in E. Scale bar, 100 μm. (E) Representative images of projection into mMN of control mice (E_1_). Bottom panel shows axons without DAPI counterstain (E_2_). Labeling in lMN are unrelated axons from pre-subiculum. Scale bar, 100 μm. (F, H, J) Same as D, but for genotypes indicated above. (G_1,2_, I_1,2_, K_1,2_) Same as E, but for genotypes indicated above. dSub axons in *Ten3^mMN^&Lphn2^mMN^*-cKO (G) and *Lphn2^mMN^-*cKO (I) mice spread medially outside of lat-mMN (arrows in G_2_, I_2_). (G_3_, I_3_, K_3_) Fraction of total projection intensity of the dSub axons in med-mMN for genotypes indicated above. n = 5, 8 for control and *Ten3^mMN^&Lphn2^mMN^*-cKO mice, respectively (G_2_); n = 6, 4 for control and *Lphn2^mMN^*-cKO mice, respectively (I_2_); n = 7, 4 for control and *Ten3^mMN^*-cKO mice, respectively (K_2_). Open circles indicate representative animals shown in E, G, I, K. Mean ± SEM. Mann-Whitney test, ** p < 0.01. (L) Summary of target selection of dSub axons in mMN in control (L_1_) or when *Ten3* (L_4_), *Lphn2* (L_3_), or both (L_2_) were conditionally knocked out in mMN. See Figures S3 and S6 for additional data.

We injected *AAV-mCh* into dSub to assess dSub→lat-mMN axon projections and, again, started by assessing *Sim1-Cre;Ten3^fl/fl^;Lphn2^fl/fl^* (*Ten3^mMN^&Lphn2^mMN^-*cKO) animals to determine if the Ten3 axonal deletion phenotype is recapitulated by *Ten3* and *Lphn2* double cKO in the target (**Figure 5B, C**). In *Ten3^mMN^&Lphn2^mMN^-*cKO animals, we observed mistargeting of the Ten3^+^ dSub axons into the Lphn2-high med-mMN when compared to *Ten3^fl/fl^;Lphn2^fl/fl^*control (**Figure 5D–G**). This mistargeting was also observed in contralateral mMN of *Ten3^mMN^&Lphn2^mMN^-*cKO (**Figure S6A, B, F**). Thus, Ten3 and/or Lphn2 is required in mMN for the appropriate targeting of Ten3^+^ dSub axons.

To determine the relative contribution of Lphn2 and Ten3 to the double target knockout phenotype, we repeated these experiments with *Lphn2* or *Ten3* single cKOs in the mMN target. Similar to the *Ten3^mMN^&Lphn2^mMN^-*cKO, Ten3^+^ dSub axons mistargeted to med-mMN in *Lphn2^mMN^*-cKO animals as well, causing an increase in the density of axons in med-mMN relative to the appropriate lat-mMN target (**Figure 5H, I**). However, dSub axons in *Ten3^mMN^-*cKO still predominantly targeted lat-mMN (**Figure 5J, K**). These phenotypes were recapitulated in contralateral mMN with mistargeting in *Lphn2^mMN^*-cKO, but not *Ten3^mMN^-*cKO animals (**Figure S6C, D**).

Taken together, these results suggest that Lphn2–Ten3-mediated repulsion is required in the dSub→lat-mMN connection (**Figure 5L**, **Figure S6E**). Additionally, the mistargeting phenotype in *Lphn2^mMN^-*cKO animals implied that, like in pCA1→dSub, Ten3-mediated attraction alone cannot compensate for the loss of Lphn2-mediated repulsion for the appropriate targeting of Ten3^+^ axons.

### Lphn2 is required in subiculum for precise targeting of pSub axons to med-mMN

So far, we have focused on connections between target selection of Ten3^+^ axons in the medial subdivision of the extended hippocampal network (**Figure 1A**). However, in the dCA1→pSub (the lateral subfields of the CA1→subiculum connection), Lphn2^+^ CA1 axons are repelled by Ten3 in dSub, meaning Ten3–Lphn2 heterophilic repulsions are reciprocal, with both Ten3 and Lphn2 capable of acting as receptor and ligand (**Figure 1A_3_**).^9^ Therefore, we asked whether reciprocal repulsions are also generalizable to other nodes in the extended hippocampal network. We used pSub→med-mMN, the Lphn2-high subdivision of the subiculum→mMN connection, to address this question.

Modifying the experiments in Figure 4 for examining Lphn2^+^ axons, we injected *AAV-CreON-GFP* into pSub of *Nts-Cre;Lphn2^+/+^* controls and *Nts-Cre;Lphn2^fl/fl^* (*Lphn2^Sub^*-cKO) mice (**Figure 6A, B**). In control animals, pSub axons were restricted to the med-mMN whereas aberrant pSub axons strayed into lat-mMN in *Lphn2^Sub^*-cKO (**Figure 6C–F**). Although this phenotype was more subtle than when *Ten3* was deleted in dSub axons, there was a significant increase of axon density in Ten3-high lat-mMN in *Lphn2^Sub^*-cKO compared to control (**Figure 6G**). This phenotype was also present in a smaller subset of contralateral mMN (**Figure 6I**). These data demonstrate that Lphn2 is required in subiculum for pSub axons to precisely target med-mMN (**Figure 6H**).

**Figure 6.**
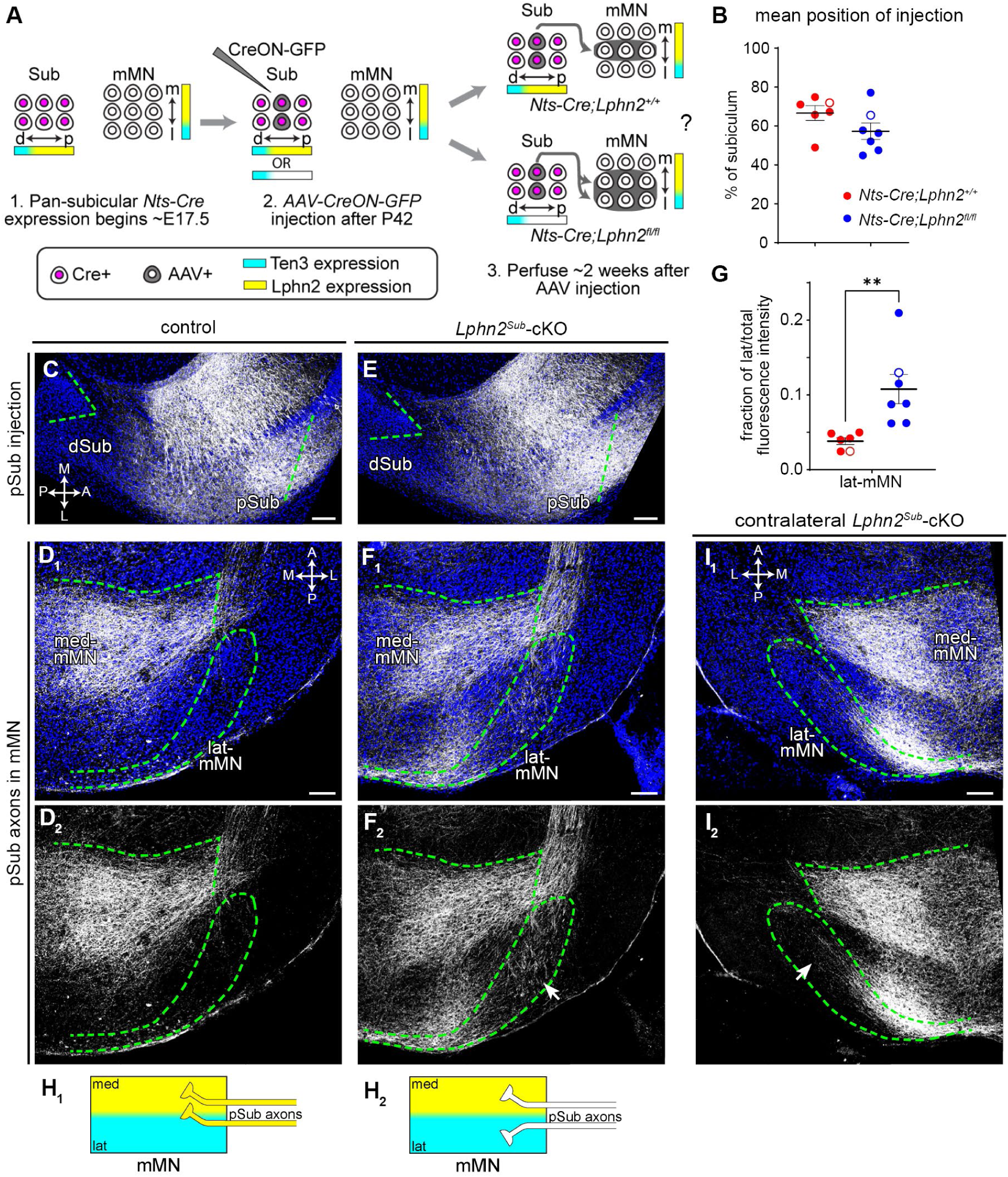
Lphn2 is required in subiculum for pSub axons to precisely target to medial mMN. (A) Injection strategy and possible results for tracing pSub axons into mMN of control (top right) and *Lphn2^Sub^-*cKO mice (bottom right). (B) Mean positions of injection sites along the dSub-to-pSub axis show no significant differences between controls (n = 6) and *Lphn2^Sub^-*cKO (n = 7) mice. Open circles indicate representative animals shown in C, E. Mean ± SEM. Mann-Whitney test. (C) Representative image of the *AAV-GFP* (gray) injection site of controls show that the virus is predominantly restricted to pSub. Same animal as in D. (D) Representative images of projection of pSub axons (gray) into mMN of controls (D_1_). Bottom panel shows axons without DAPI counterstain (D_2_). (E, F) Same as C, D, but for *Lphn2^Sub^-*cKO mice. *Lphn2* cKO axons spread outside of med-mMN and into lat-mMN (arrow in F_2_). (G) Fraction of total projection intensity of pSub axons in lat-mMN of controls (n = 6; red) and *Lphn2^Sub^-*cKO (n = 7; blue). Open circles indicate representative animals shown in D, F. Mean ± SEM. Mann-Whitney test, ** p < 0.01. (H) Summary of target selection of pSub axons in mMN of control (H_1_) and *Lphn2^Sub^*-cKO (H_2_) mice based on results in C–G. (I) Same as F, but for projections into contralateral mMN. Contralateral *Lphn2* cKO axons spread into lat-mMN (arrow in I_2_). Scale bars, 100 μm. See Figures S3 and S7 for additional data.

### Target requirement for the pSub→med-mMN projection

To determine if Ten3–Lphn2 are required in the mMN target for the appropriate targeting of pSub→med-mMN and, thus, recapitulate the *Lphn2^Sub^*-cKO phenotype, we injected *AAV-GFP* into pSub of *Sim1-Cre;Ten3^fl/fl^;Lphn2^fl/fl^* (*Ten3^mMN^&Lphn2^mMN^-*cKO) animals (**Figure 7A, B**). We observed aberrant Lphn2^+^ pSub axons in the Ten3-high lat-mMN when compared to *Ten3^fl/fl^;Lphn2^fl/fl^*controls (**Figure 7C–G**), including in a few contralateral mMN (**Figure 7I**), recapitulating the spreading phenotype of *Lphn2* deletion in subiculum axons (**Figure 6**). Because Lphn2 does not exhibit homophilic binding,^9,47^ we did not test single cKOs as it is unlikely that loss of Lphn2 in the target contributes to the phenotype in *Ten3^mMN^&Lphn2^mMN^-*cKOs. Taken together, these data suggest that Ten3–Lphn2 heterophilic repulsion is required to correctly specify pSub→med-mMN (**Figure 7H**) and that this repulsion is reciprocal within the subiculum→mMN connection, as in the CA1→subiculum connection^9^ (**Figure 7J**).

**Figure 7.**
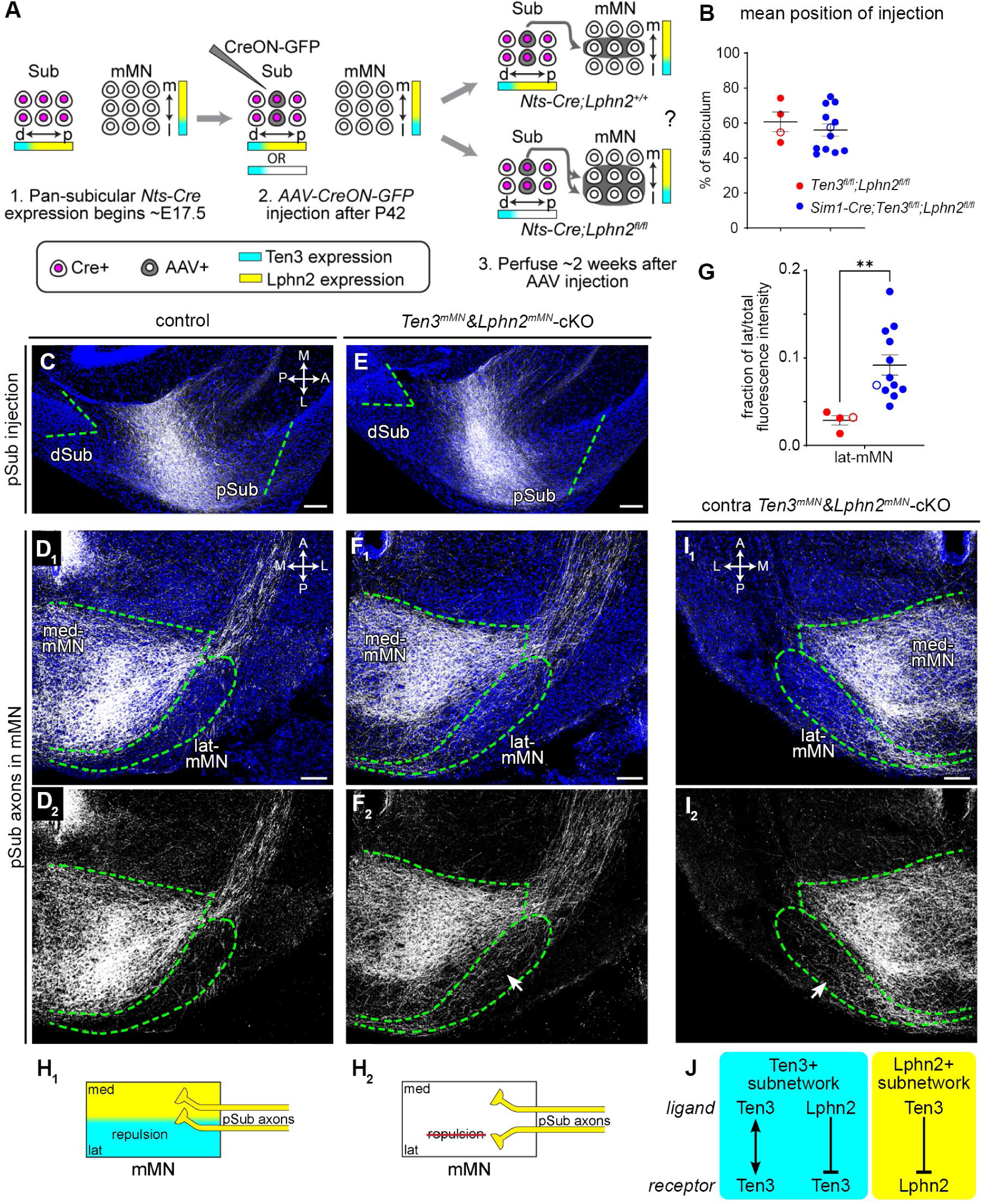
Ten3/Lphn2 are required in mMN for the precise targeting of pSub axons. (A) Injection strategy and possible results for tracing pSub axons into mMN of control (top right) and *Ten3^mMN^&Lphn2^mMN^-*cKO mice (bottom right). (B) Mean positions of injection sites along the dSub-to-pSub axis show no significant differences between controls (n = 4, red) and *Ten3^mMN^&Lphn2^mMN^-*cKO (n = 12, blue). Open circles indicate representative animals shown in C, E. Mean ± SEM. Mann-Whitney test. (C) Representative image of the *AAV-GFP* (gray) injection site of controls show that the virus is predominantly restricted to pSub. Injection corresponds to animal in (D). (D) Representative images of projection of pSub axons (gray) into mMN of control mice (D_1_). Bottom panel shows axons without DAPI counterstain (D_2_). (E, F) Same as C, D, but for *Ten3^mMN^&Lphn2^mMN^-*cKO mice. Lphn2^+^ pSub axons spread into lat-mMN (arrow in F_2_). (G) Fraction of total projection intensity of pSub axons in lat-mMN of controls (n = 4 mice, red) and *Ten3^mMN^&Lphn2^mMN^-*cKO (n = 12 mice, blue). Open circles indicate representative animals shown in D, F. Mean ± SEM. Mann-Whitney test, ** p < 0.01. (H) Summary of target selection of pSub axons in mMN of control (H_1_) and *Ten3^mMN^&Lphn2^mMN^*-cKO (H_2_) mice based on results in C–G. (I) Same as F, but for projections into contralateral mMN. Lphn2^+^ pSub axons spread into contralateral lat-mMN of *Ten3^mMN^&Lphn2^mMN^*-cKO mMN (arrow in I_2_). (J) Summary of the Ten3–Lphn2 molecular module. Originally established in a CA1→Sub projection, this study demonstrates its broad use across the extended hippocampal network. Scale bars, 100 μm. See Figure S3 for additional data.

### Lack of evidence that Lphn2–Ten3 mediates axon-axon interaction

Ligand–receptor pairs could mediate axon-axon interactions in addition to axon-target interaction.^26,48^ Given that both Ten3^+^ and Lphn2^+^ axons from subiculum can serve as repulsive receptors for their respective ligand in mMN, we asked whether these molecules could also mediate axon-axon repulsion, which could, in principle, sort Ten3^+^ dSub axons and Lphn2^+^ pSub axons prior to reaching the mMN target. This target-independent sorting could contribute to the precise target selection of these two populations of axons. To test this, we traced wildtype Lphn2^+^ axons from pSub into their mMN target in *Ten3^Sub^*-cKO animals to determine if loss of Ten3 on neighboring axons affected their targeting **(Figure S7A, F).** We did not observe a significant difference in control and *Ten3^Sub^*-cKO animals; in both cases, Lphn2^+^ pSub axons targeted to med-mMN appropriately, with no aberrant axons in lat-mMN **(Figure S7B–E, G).** We also performed the converse experiment where we deleted *Lphn2* in subiculum (*Lphn2^Sub^*^-^cKO) and traced wildtype Ten3^+^ axons from dSub into their mMN target to determine if loss of Lphn2 on neighboring axons affected their targeting **(Figure S7I, N).** Again, we did not detect a significant difference in targeting between *Lphn2^Sub^*-cKO and control mice, indicating little to no target-independent axon-axon interaction **(Figure S7J–M, O).** Taken together, these data argue against direct interaction between dSub and pSub axons playing a role in target selection at mMN (**Figure S7H, P**). These results are consistent with our finding that dSub and pSub axons were intermingled prior to arrival at mMN, both in developing and adult brains **(Figure S1B, F, G)** and that expression onset of Ten3 and Lphn2 seems to coincide with axons arrival in the targets but not before **(Figure S1D**).^8,39^ These data also provide strong support that the phenotypes we observed in **Figures 4–7** are due to interactions occurring exclusively in the target.

## DISCUSSION

Our systematic conditional knockout approach demonstrated that Ten3–Ten3 homophilic attraction and Ten3–Lphn2 heterophilic reciprocal repulsion (**Figure 7J**) instruct target selection within the extended hippocampal network—MEC→dSub/pCA1, dSub→lat-mMN, and pSub→med-mMN, in addition to previously reported CA1→subiculum connections.^8,9^ To our knowledge, this is the first time a ligand–receptor pair has been shown to instruct target selection in multiple nodes of the same network independently. Our analysis of conditional target deletions also revealed differential requirements of repulsion and attraction in different nodes possibly depending on the order in which developing axons encounter the attractant versus the repellent subfield in the target (**Figure 1F**).

### Differential requirement of attraction and repulsion at different circuit nodes

In the pCA1→dSub connection, our previous studies indicated that target deletion of *Lphn2* alone caused spread of pCA1 axons to pSub, and target deletion of both *Lphn2* and *Ten3* exhibited more severe pCA1 axon spreading to pSub.^9^ Thus, Lphn2–Ten3 repulsion is required for preventing pCA1 axon mistargeting. This is likely because Ten3^+^ pCA1 axons first encounter the Lphn2^+^ pSub subfield before Ten3^+^ dSub subfield (**Figure 1F, left**), making repulsion necessary to prevent axons stalling before they find the attractive subfield.

By contrast, conditional target deletion of *Lphn*2 did not cause a significant targeting defect in the MEC→dSub connection (**Figure 3I**). Only when we deleted both *Ten3* and *Lphn2* in the subiculum target did we recapitulate *Ten3* deletion in MEC axon (**Figure 3G**, compared to **Figure 2H**). (Subiculum deletion of *Ten3* and *Lphn2* has a weaker phenotype than *Ten3* deletion in MEC axons possibly due to the late onset of *Nts-Cre,* between E17.5 and P0, such that some target proteins may have already been produced before conditional gene knockout.) Our developmental axon tracing indicated that MEC axons first encountered Ten3^+^ dSub before encountering Lphn2^+^ pSub (**Figure 1F, middle**; **Figure S1D, E**). Thus, the simplest interpretation for the lack of phenotype of *Lphn2* target deletion in the MEC→dSub connection is that Ten3–Ten3 homophilic attraction is largely sufficient to retain MEC axons in the Ten3^+^ dSub subfield. Repulsion can increase the robustness of the system—to ensure correct wiring even when attraction is disrupted.

Interestingly, when Ten3^+^ axons simultaneously encounter the Ten3^+^ and Lphn2^+^ target subfields, as is the case for the dSub→lat-mMN connection (**Figure 1F, right**; **Figure S1F, G**), conditional target deletion of *Lphn2* alone caused a robust mistargeting phenotype of Ten3^+^ dSub axons entering the Lphn2^+^ med-mMN subfield (**Figure 5I**). This indicates that Ten3–Ten3 attraction cannot compensate for the loss of Lphn2–Ten3 repulsion and highlights the importance of simultaneous push and pull in ensuring the correct axon sorting at the choice point. Altogether, these comparative analyses highlight the variations on the basic theme of cooperation between attraction and repulsion during target selection.

Incidentally, the order in which axons encounter the attractant versus repellent subfields also appears to influence the severity of mistargeting when it occurs. In *Ten3* and *Lphn2* double conditional target knockout, the MEC→dSub axons mistarget only to adjacent Lphn2-high regions (**Figure 3**), whereas dSub→lat-mMN axons can mistarget to the most medial region of med-mMN (**Figure S5**). Although time-lapse studies would be ideal to confirm this proposal, these data suggest that the order in which axons encounter attractant and repellent subfields in the target is an important factor to consider in developmental studies.

### Multi-functionality and repeated use highlight the economy of wiring molecules

Our study highlights two mechanisms that enable the developing brain to use a limited number of cell-surface recognition proteins to wire up many more neurons and connections. First, wiring molecules appear to be highly multifunctional, with both Ten3 and Lphn2 serving as both ligand and receptor. Even within the same neuron, Ten3 can simultaneously mediate both attraction and repulsion depending on its interacting partners. For example, in dSub neurons, Ten3 acts cell non-autonomously as an attractive ligand for MEC axons (**Figure 3**) and pCA1 axons,^8^ and as a repulsive ligand for dCA1 axons^9^. At the same time, it acts cell autonomously as a receptor for the Lphn2 repellent in mMN as dSub axons select the appropriate subfield to make connections (**Figure 4**). Similarly, Lphn2 in pSub neurons acts cell non-autonomously as a repulsive ligand for pCA1 axons^9^ and for MEC axons (**Figure 3**), while acting cell autonomously as a receptor for target selection in mMN (**Figure 6**). We note that members of the teneurin and latrophilin families also function to regulate synapse formation in later stages of development within these same regions.^40,49,50^

Second, the same molecular module—Ten3–Ten3-mediated attraction and Ten3–Lphn2-mediated mutual repulsions (**Figure 7J**)—appears to be repeatedly used across multiple nodes of the same circuit and potentially many circuits across the brain. We have used systematic conditional knockouts to demonstrate conclusively the repeated use of this molecular module in instructing connections in the extended hippocampal network. Our companion manuscripts reported the broad deployment of inverse expression of Ten3 and Lphn2 across many additional circuits in the mouse brain and the spinal cord.^39,51^ In all tested cases, Ten3–Lphn2 inverse expression patterns lay out a topographic map in the developing brain prior to the establishment of functional connections. Importantly, to ensure Ten3^+^/Lphn2^+^ neurons from one node do not encounter Ten3^+^/Lphn2^+^ neurons from another circuit to produce unintended cross interactions, spatiotemporal control of their expression and the collaboration with other guidance molecules are likely necessary.

The multi-functionality and repeated use of the Ten3–Lphn2 module are both a result of the molecular property of Ten3 and Lphn2 proteins as a ligand–receptor pair (**Figure 7J**) and the circuit architecture of the extended hippocampal network, featuring convergent, divergent, and recurrent connectivity motifs (**Figure 1A_2_**).^52^ Although many proteins are multifunctional, Ten3 in subiculum neurons serves dual functions in two compartments of the same cell, which is of particular benefit for reducing number of wiring molecules in highly interconnected regions. Given the prevalence of these connectivity motifs in the mammalian central nervous system, we envision that such multi-functionality of wiring molecules will become a theme when detailed functional analyses are carried out on other molecules and in other circuits. Finally, we note that conditional knockout of *Lphn2* in subiculum causes a shift in place cell distribution consistent with our miswiring phenotypes,^53^ supporting the functional importance of topographic mapping mediated by the Lphn2–Ten3 module.

### Comparisons with ephrin-A–EphA and cadherins

Classic work has shown that ephrinA–EphA mediate anterior-posterior retinotopic target selection in retina→SC and, to a lesser extent, retina→LGN and LGN→V1.^5,6,19,23–25^ However, since ephrinAs were removed from all targets and axons simultaneously, it is difficult to distinguish whether ephrinAs act as ligands or receptors, and whether the phenotype observed in one circuit node is caused by direct action of ephrinAs within that node or is a secondary consequence of miswiring in an upstream node. By using axon- and target-specific knockouts of both Ten3 and Lphn2, we demonstrate conclusively that the Lphn2–Ten3 module directly regulates target selection within each node. Additionally, while ephrin-A2, A3, and A5 serve largely redundant functions in specifying topography, only one family member of receptor and ligand seems sufficient in the Lphn2–Ten3 module. Finally, compared to the well-investigated ephrin–Eph and cadherin systems for target selection, which predominantly operate through reciprocal repulsion and homophilic attraction respectively,^10,19,54^ the Lphn2–Ten3 module incorporates both features. Cooperative push-pull of the Lphn2–Ten3 heterophilic repulsion and Ten3–Ten3 homophilic attraction can increase the robustness of this molecular module, potentially contributing to its wide deployment^39,51^ beyond the extended hippocampal network investigated here.

## RESOURCE AVAILABILITY

### Lead contact

Further information and requests for resources should be directed to the lead contact, Liqun Luo.

### Materials availability

This study did not generate new unique reagents.

### Data and code availability

- Data reported in this paper are available from lead contact upon request.
- Data was normalized using custom MATLAB code (resample.m), previously published in Pederick et al. 2021^9^ and can be found here: https://github.com/dpederick/Reciprocal-repulsions-instruct-the-precise-assembly-of-parallel-hippocampal-networks
- Any additional information required to reanalyze the data reported in this paper is available from the lead contact upon request.

## ACKNOWLEDGEMENTS

We thank the Neuroscience Gene Vector and Virus Core at Stanford University and SignaGen for producing and packaging custom viruses, T. Südhof for the *Lphn2^mVenus^*and *Lphn2^floxed^* mice, and members of the Luo lab, especially U. Chon, for advice and support. We thank T. Hindmarsh-Sten, A. Starr, Z. Li, K. Sangster, and A. Kania for critiques of the manuscript. E.C.G. was supported by National Institute of Mental Health, Ruth L. Kirschstein National Research Service Award (F31MH129079). D.T.P. was supported by an American Australian Association Education Fund Scholarship and the SFARI Fellows-to-Faculty Award (Simons Foundation). L.L. is an investigator at Howard Hughes Medical Institute. This work was supported by National Institutes of Health grant (R01-NS050580 to L.L.).

## AUTHOR CONTRIBUTIONS

E.C.G. and D.T.P. conceptualized the study. E.C.G. performed all the experiments, except for RNAscope, performed by Y.Z. E.C.G. analyzed all data and wrote the manuscript. L.L. supervised the study and edited the manuscript.

## DECLARATION OF INTERESTS

The authors declare no conflicts of interest.

## STAR METHODS

### Key Resource Table

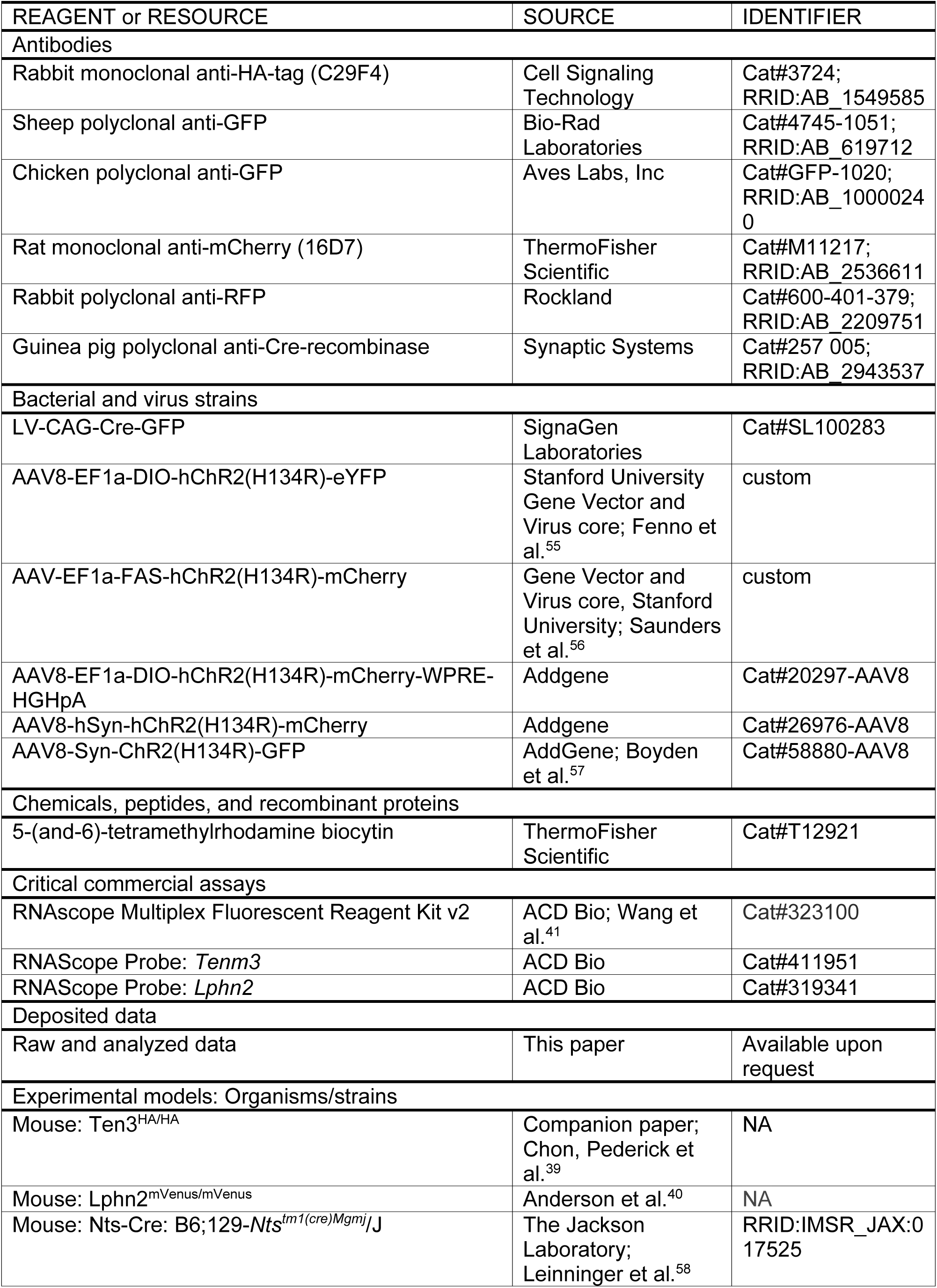
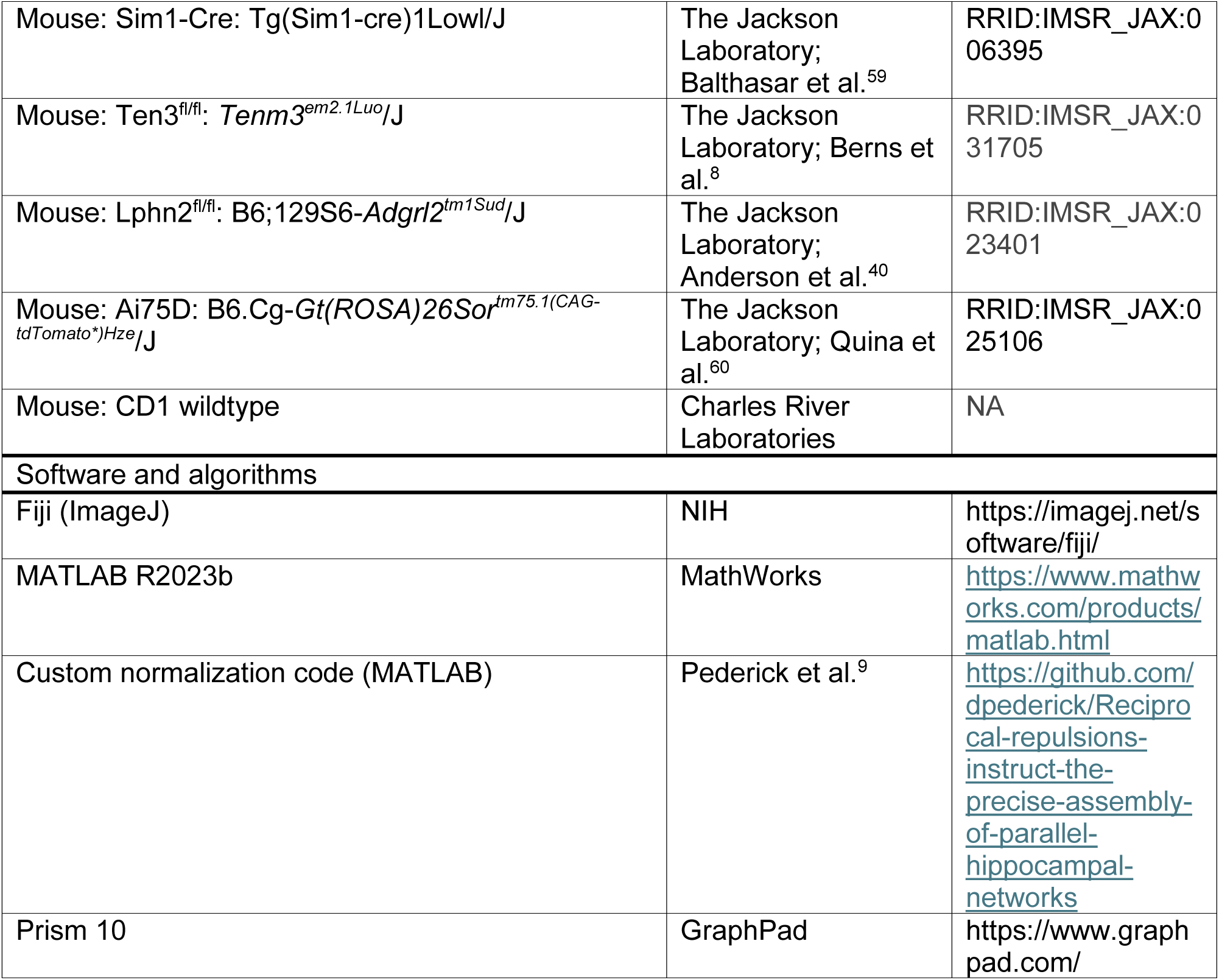

## EXPERIMENTAL MODEL AND STUDY PARTICIPANTS

Protein expression and *ex vivo* injection experiments used *Ten3^HA/HA^;Lphn2^mVenus/mVenus^*mice at postnatal day 0 (P0) and P8.^39,40^ CD-1 mice from Charles River Laboratories were used for double *in situ* hybridization and *ex vivo* tracing timeline experiments, ages ranging from embryonic day 15.5 (E15.5) to P6. All loss-of-function (LOF) experiments used crosses of *Nts-Cre* (C57BL/6 and 129/SvJ mixed background; JAX #017525),^58^ *Sim1-Cre* (C57BL/6, 129/SvJ, and FVB mixed background; JAX #006395),^59^ *Ten3^fl/fl^* (mixed background C57BL/6, 129/SvJ, and CD1),^8^ and *Lphn2^fl/fl^* (mixed background C57BL/6, 129/SvJ, and CD1).^40^ Cre lines were characterized at E15.5, E17.5, P0 and adult using Ai75D nuclear reporter mice (*Rosa-CAG-LSL-ntdTomato::deltaNeo*; C57BL/6; JAX stock #025106).^60^ Genotypes used for each experiment are listed in detail in Table S1.

For the entorhinal cortex→subiculum and entorhinal cortex→CA1 *Ten3* deletion in axons (CreON/CreOFF) experiments (Figure 2, Figure S2), breeder pairs were maintained to produce all homozygous litters due to the low probability of correctly injecting overlapped P0 lentivirus and adult AAVs. Thus, these animals were not randomly allocated to experimental groups and were instead compared to age-matched CD1 wildtypes. Randomly allocated littermate controls were used for all other loss-of-function experiments conducted at P42 or later.

Due to low efficiency and strict cut off criteria, the total number of mice injected and screened for each experiment is as follows: Figure 2/Figure S2: 161 *Ten3^+/+^* and 181 *Ten3^fl/fl^*; Figure 3/Figure S4: 73 *Nts-Cre;Ten3^fl/fl^;Lphn2^fl/fl^*and controls, 136 *Nts-Cre;Lphn2^fl/fl^* and controls, and 43 *Nts-Cre;Ten3^fl/fl^* and controls; Figure 4/Figure S5, S7: 146 *Nts-Cre;Ten3^fl/fl^*and controls; Figure 5, 7/Figure S6: 62 *Sim1-Cre;Ten3^fl/fl^;Lphn2^fl/fl^*and controls, 67 *Sim1-Cre;Lphn2^fl/fl^* and controls, and 55 *Sim1-Cre;Ten3^fl/fl^* and controls; Figure 6/Figure S7: 66 *Nts-Cre;Lphn2^fl/fl^*and controls.

Male and female mice were used for all experiments. Mice were group-housed on a 12 hr light/dark cycle with *ad libitum* access to food and water. All animal and virus procedures followed guidelines approved by Stanford University’s Administrative Panel on Laboratory Animal Care and Institutional Biosafety Committee in accordance with NIH guidelines.

## METHOD DETAILS

### Immunostaining

Animals were anesthetized with 2.5% Avertin (intraperitoneal) and perfused (transcardial) with 10 mM PBS, followed by 4% paraformaldehyde (PFA). Brains were dissected and post-fixed in 4% PFA for 1–2 hours (P0, P8) or 4–6 hours (>P42), then cryo-protected for at least 48 hours in a 30% sucrose solution. E15.5 and E17.5 brains for Cre characterizations (Figure S3) were fresh-dissected and post-fixed in 4% PFA overnight before cryoprotection. Brains were embedded in Optimum Cutting Temperature (OCT)-filled cryomolds in a 2-methylbutane bath, cooled by dry ice and stored at –80°C until sectioning. Serial 60-μm horizontal sections were collected free-floating in 24-well plates in a solution of 10 mM PBS and 0.02% sodium azide and stored at 4°C until staining. CD1 wildtype *ex vivo* injections were sectioned at 180-μm (see ‘*Ex vivo* injections’ for more details). The staining procedure was as follows: Incubation in (1) 10 mM PBS for 3’ 10 min washes at room temperature (RT), (2) blocking solution of 0.3% PBST and 10% normal donkey serum (NDS) for 1–2 hours RT, (3) primary antibody in 0.3% PBST for 2–3 nights at 4°C, (4) 0.3% PBST for 3’ 10 min washes RT, (5) secondary antibody in 0.3% PBST and 10% NDS block for 1–2 hours RT, (6) DAPI (1:2500 dilution of 5 mg/mL; Invitrogen) in 0.3% PBST for 15 min RT, and (7) 10 mM PBS for 3’ 10 min washes RT. Sections were mounted in 10 mM PBS and cover-slipped using ProLong Gold Antifade Mountant (Invitrogen). Primary antibodies include: Rabbit anti-HA (1:300, Cell Signaling #3724), Sheep anti-GFP (1:1500, BioRad 4745-1051), chicken anti-GFP (1:1000, Aves Labs, GFP-1020), rat anti-mCherry (1:1000, ThermoFisher, M11217), rabbit anti-RFP (1:1000, Rockland 600-401-379), and guinea pig anti-Cre-recombinase (1:500, Synaptic Systems 257 005). Secondary antibodies used were species-specific conjugated antibodies: AlexaFluor 488 (1:1000, Invitrogen), Cy3 Affinipure (1:1000, Jackson ImmunoResearch), AlexaFluor 647 (1:1000, Invitrogen). Images were taken with a Zeiss LSM 900 confocal microscope. For quantification details see ‘Image and data analysis’ sections.

### Double *in situ* hybridization (RNAScope)

E15.5, E17.5, and P0 pups were sacrificed by rapid decapitation. For embryonic dissections, pregnant dams were anesthetized with isoflurane before cervical dislocation and uterine dissection. Brains were fresh-dissected in ice-cold 10 mM PBS and immediately embedded and frozen in OCT-filled cryomolds in a 2-methylbutane bath, cooled by dry ice. Brains were stored at –80°C until sectioning. Serial 12-μm horizontal sections were collected on Superfrost Plus slides (Fisher Scientific) and stored at –20°C until procedure. Double *in situs* were performed using the RNAscope Multiplex Fluorescent Reagent Kit v2 (ACD Bio)^41^ and performed according to RNAScope assay guidelines for fresh-frozen samples with the following modifications for neonatal tissue during pretreatment: Sections were fixed on slides for 90 minutes in 4% PFA and Protease III was used during the digestion step. Probes used were RNAScope Probe-Mm-Tenm3 (1:2, ACD Bio 411951) and RNAScope Probe-Mm-Lphn2-C2 (1:100, ACD Bio 319341-C2). Fluorophores used were Opal Dye 520, 570, and 690 (1:1500, Akoya Biosciences). Images were taken with a Zeiss LSM 900 confocal microscope (10’ objective, with 0.5’ digital zoom).

### *Ex vivo* injections

For embryonic dissections, pregnant dams were anesthetized with isoflurane before cervical dislocation and uterine dissection. Pups were sacrificed by rapid decapitation individually at the time of injection. Brains were fresh-dissected into ice-cold artificial cerebrospinal fluid (ACSF) that contained (in mM) 110 choline chloride, 2.5 KCl, 1.2 NaH_2_PO_4_, 20 HEPES, 5 sodium ascorbate, 2 thiourea, 3 sodium pyruvate, 1 CaCl_2_, 1.3 MgCl_2_, 26.8 NaHCO_3_, 25 glucose, and 73 trehalose. A scalpel was used to crudely remove olfactory bulbs and the most rostral ∼1/4 of the brain to create a flat surface as well as any midbrain and hindbrain structures obscuring the caudal surface of cerebral cortex. Brains were pinned with caudal surface exposed in a silicon-lined dissecting dish filled with ice-cold ACSF. Using exterior anatomy as a guide, 5% 5-(and-6)-tetramethylrhodamine biocytin in 0.9% saline (biocytin TMR; Invitrogen) was iontophoretically injected into the approximate location of medial entorhinal cortex. Iontophoresis was performed for 3 min at 5 μA current using glass pipette tips with an outside diameter of 15–20-μm. Brains were then placed in carbogenated ACSF at RT for ∼6 hours, covered to protect from light, to allow time for dye to spread to axon terminals. After 6 hours, brains were placed in 4% PFA overnight to fix and moved to 30% sucrose solution for 24–48 hours for cryoprotection. Brains were embedded, sectioned, and stained as described previously (see ‘Immunostaining’), but were sectioned at 180-μm for axon tracing in wildtype animals to visualize more of the axon trajectory in one plane (Figure S1C, D).

### Stereotactic injections in neonatal pups

Pups were anesthetized via hypothermia and mounted in stereotactic apparatus modified to hold them. Cranial anatomy was visualized through the skin for stereotactic coordinates and a small incision was created using a 27-gauge needle in place of a craniotomy. Stereotactic coordinates were measured from lambda. MEC coordinates were 2.05–2.15 mm lateral, 0.6–0.65 mm posterior, and 1.4–1.45 mm ventral from brain surface. 400 nL of *LV-CAG-Cre-GFP* (∼6.6x10^8^ copies/mL; SignaGen SL100283) was injected at 100 nl/min.

### Stereotactic injections in adult mice

P42 or older mice were anesthetized using isoflurane and mounted in stereotactic apparatus. Virus was injected via iontophoresis (5 uA current, 7 sec on/7 sec off) for 2 min using glass pipette tips with an outside diameter of 10–15-μm. Mice were perfused 2–3 weeks post-injection and processed as described above (see ‘Immunostaining’). Stereotactic coordinates were all measured from lambda and vary based on strain, genotype, and age. MEC coordinates were 2.75–3.1 mm lateral, 0.475–0.675 mm posterior, and 2.05–2.2 mm ventral from brain surface. Distal subiculum (dSub) coordinates were 2.8–2.9 mm lateral, 0.175 mm posterior to 0.05 mm anterior, and 1.95–2.05 mm ventral from brain surface. Proximal subiculum (pSub) coordinates were 3.05–3.1 mm lateral, 0.2–0.525 mm anterior, and 1.95–2.05 mm ventral from brain surface. Viruses injected were as follows: *AAV8-EF1a-DIO-hChR2(H134R)-eYFP* (*CreON-GFP*; 2 x10^12^–1x10^13^ copies/mL; Gene Vector and Virus core, Stanford University)^55^, *AAV8-EF1a-FAS-hChR2(H134R)-mCherry* (*CreOFF-mCh*; 2–8x10^12^ copies/mL; Gene Vector and Virus core, Stanford University),^56^ *AAV8-EF1a-DIO-hChR2(H134R)-mCherry-WPRE-HGHpA* (*CreON-mCh*; 4x10^12^–1.2x10^13^ copies/mL; gift from Karl Deisseroth, Addgene viral prep # 20297-AAV8), *AAV8-hSyn-hChR2(H134R)-mCherry* (*AAV-mCh*; 1x10^13^ copies/mL gift from Karl Deisseroth, Addgene viral prep # 26976 -AAV8), and *AAV8-Syn-ChR2(H134R)-GFP* (*AAV-GFP*; 1x10^13^ copies/mL; gift from Edward Boyden, Addgene viral prep # 58880-AAV8).^57^

## QUANTIFICATION AND STATISTICAL ANALYSIS

### Image and data analysis for entorhinal cortex→subiculum axon tracing

Images of injection sites (10’ magnification) and projections (10’ objective, with 0.45’ digital zoom) were acquired for every other 60-μm section using a Zeiss LSM 900 confocal. Images of direction of axons entering subiculum at P0 were taken at 20’ with 0.5’ digital zoom (Figure S1E). Due to variations in injection sites and time lapse between experiments, exposures were adjusted for each mouse and fluorescence intensity was normalized post hoc for comparison. Fluorescence intensity and injection position measurements were performed on unprocessed images and collected using FIJI. Data processing was performed in MATLAB and Excel.

Strict inclusion criteria were applied to ensure tracing only from the appropriate subnetwork (Ten3^+^ MEC). Inclusion criteria were as follows: (1) Both AAV injection sites must be in medial entorhinal cortex (mean position within most medial 20% of entorhinal cortex)—based on Ten3 mRNA in horizontal entorhinal section^9^; (2) lentivirus *Cre-GFP* injection must allow for roughly equal expression of *AAV-CreON-GFP* and *AAV-CreOFF-mCh;* (3) 3 brightest sections of *AAV-CreON-GFP* and *AAV-CreOFF-mCh* must overlap by 2 of 3 sections across the dorsal–ventral axis; (4) lentivirus Cre-GFP expression in subiculum and CA1 must be minimal. Animals that fulfilled these criteria were included in the quantifications reported in Figures 2, 3 and Figures S2, 4.

For injection site quantification, all image and data processing were done using FIJI. First, a 425-pixel-wide segmented line was drawn from medial to lateral of layer II/III entorhinal cortex using DAPI counterstain to define edges. The segmented lines from each channel in the image were straightened using the Straighten function. Background subtraction was done by measuring the mode value of background regions and using the Subtract function to remove the mode value. The Plot Profile command was then used to extract fluorescence intensity (FI) along the medial–lateral axis. To compare across animals, these FI plots were resampled into 100 equal bins representing the length of entorhinal cortex using a custom MATLAB code.^8,9^ The FI traces of the three brightest sections of injection site were summed together and normalized to FI values of 0–100. The mean position of the injection site was calculated by multiplying the FI value by its bin position, summing across the entire axis, and dividing by the sum of FI values.

Because *LV-Cre-GFP* may extend past the *AAV-CreON-GFP* tracing injection, it contributes fluorescence that would influence the calculation for AAV injection mean position. To resolve this, all animals were stained with a Cre antibody in the far-red channel that was also processed as above. The image calculator function was used to subtract the Cre only channel from the GFP channel to remove any GFP being contributed by Cre-GFP before the Plot Profile was extracted.

Projection site processing and quantification was done similarly to the injection site quantification, but the initial line was drawn as a 50-pixel wide segmented line along the distal–proximal axis of subiculum molecular layer and a 100-pixel-wide segmented line along the distal–proximal axis of CA1 molecular layer. Although it is impossible to avoid including axons occasionally traversing the space between subiculum and CA1, they are relatively uniform across genotype and account for only a small fraction of FI. Areas under the curve for subiculum FI traces were calculated in GraphPad Prism 10.

### Image and data analysis for subiculum→mMN expression and topography

All images were acquired for every other 60-μm section using a Zeiss LSM 900 confocal (subiculum/mMN P8 Ten3–Lphn2 expression and adult subiculum injection site (10’ objective, with 0.5’ digital zoom); adult mMN projection site (10’ objective, with 0.45’ digital zoom). Hippocampal-entorhinal image shown (Figure 1B) is at 10’ magnification. Images of axon tract and direction of axons entering mMN at P0 were taken at 20’ with 0.5’ digital zoom (Figure S1F, G).

For subiculum P8 Ten3–Lphn2 expression quantification, a 150-pixel-wide segmented line was drawn along the distal–proximal axis of the subiculum cell body layer using DAPI counterstain to define edges. The segmented line was straightened, background subtracted, quantified, and the data processed as described for MEC injection above (see ‘Image and data analysis for subiculum→mMN axon tracing’). Six sections from one animal (Figure 1B_1_) were averaged for Ten3 and Lphn2 fluorescence intensity traces (Figure S1A_1_). For mMN P8 Ten3–Lphn2 expression quantification, a 250-pixel-wide segmented line was drawn along the Ten3–Lphn2 gradient (posterolateral to anteromedial) and processed in the same way. Three sections from one animal (Figure 1B_2_) were averaged for Ten3 and Lphn2 fluorescence intensity (Figure S1A_2_).

In FIJI, a 125-pixel-wide segmented line was drawn for adult subiculum injection site, and a 250-pixel-wide segmented line was drawn for adult mMN projection site quantification. Both were processed as above, but a heatmap was generated in GraphPad Prism 10 for opposing mCherry (dSub) and GFP (pSub) injection (Figure S1A_1_) and projection site intensity (Figure S1A_2_). Three sections from one animal (Figure 1C) were averaged for each.

### Image and data analysis for subiculum→mMN axon tracing

Images of injection sites (10’ objective, with 0.5’ digital zoom) and projections (10’ objective, with 0.45’x digital zoom) were acquired for every other 60-μm section using a Zeiss LSM 900 confocal. Due to variations in injection sites and time lapse between experiments, exposures were adjusted for each mouse and fluorescence intensity was normalized post hoc for comparison. Fluorescence intensity and injection position measurements were done on unprocessed images and collected using FIJI. Data processing was performed in MATLAB and Excel.

Strict inclusion criteria were applied to ensure tracing only from the appropriate subnetwork. Inclusion criteria for Ten3^+^ dSub tracing is that AAV injection site must be in the most distal 20% of subiculum by mean position for sparse axon tracing (Figure S5) or between 12% and 20% for region-based analyses (Figures 4, 5). Animals that fulfilled these criteria were included in the quantifications reported in Figures 4, 5 and Figures S5, 6. Inclusion criteria for Lphn2^+^ pSub are as follows: AAV injection site (1) must be between 40–80% of subiculum by mean position and (2) must have less than 10% of its total fluorescence within dSub (distal 20% of subiculum). Animals that fulfilled these criteria were included in the quantifications reported in Figures 6 and 7. These criteria are based on Ten3 and Lphn2 mRNA in horizontal subiculum sections.^9^

For injection site quantification, a 125-pixel-wide segmented line was drawn along the distal–proximal axis of the subiculum cell body layer using DAPI counterstain to define edges. The segmented line was straightened, background subtracted, quantified, and the data processed as described for MEC above (see ‘Image and data analysis for subiculum→mMN axon tracing’), except without Cre-related fluorescence subtraction.

For projection site quantification, regions of interests (ROIs) of lat-mMN and med-mMN were drawn using the DAPI counterstain to define edges (Figure S5H). Background subtraction was done by measuring mean fluorescence intensity of background and using the Subtract function to delete it. Images were binarized and then the total fluorescence intensity of each ROI was collected using the raw integrated density measurement. Data is represented as the ratio of fluorescence intensity in lat-mMN (for mSub axons) or med-mMN (for dSub axons) over total fluorescence intensity in all of mMN.

### Rank order data analysis for dSub→lat-mMN axon tracing

Imaging, inclusion criteria, and image and data processing for subiculum injection site was the same as above. Imaging and image processing for mMN projection was same as above (see ‘Image and data analysis for subiculum→mMN axon tracing’). For rank order analysis, the total fluorescence intensity was collected from unprocessed images using the raw integrated density measurement in FIJI for each mMN. The two brightest ipsilateral and two brightest contralateral sections of mMN for each animal were auto-balanced using the brightness/contrast adjustment tool in FIJI to enhance the contrast of single axons. An independent researcher, blinded to genotype, numbered all ipsilateral and all contralateral images from most severe mistargeting (#1) to no mistargeting (#16), accounting for the amount of axons within the axon tract as a proxy for injection site density. Animals were re-identified and rank-sum statistics were performed (Figure S5).

### Statistics

Statistical tests were performed using GraphPad Prism 10. For sample sizes and details of statistical analyses, see figure legends. All experiments were randomized, except for the experiment of the entorhinal cortex→subiculum, *Ten3* deletion in axons (see ‘Mice’ for more details).

---

**Figure S1.**
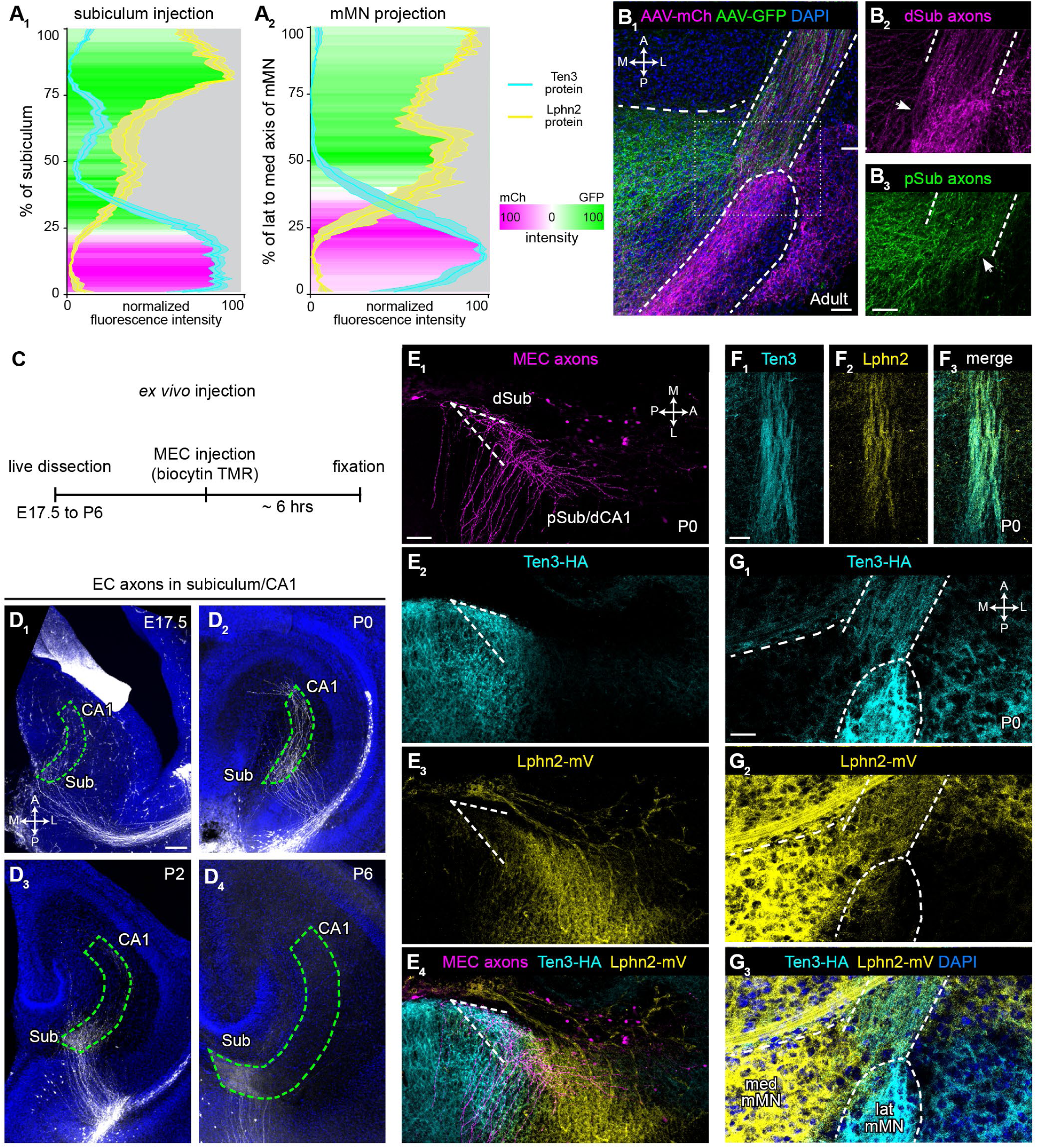
Additional data on axon tracing in adult and during development, related to Figure 1. (A) Quantitative depiction of dual-color anterograde tracing and Ten3/Lphn2 expression in subiculum→mMN projection in Figure 1B, C. (A_1_) Normalized fluorescence intensity traces of Ten3 (cyan) and Lphn2 (yellow) protein expression along the distal-to-proximal axis of subiculum in P8 *Ten3^HA/HA^;Lphn2^mVenus/mVenus^*mouse (Figure 1B_1_) overlaid with dSub (mCherry, magenta) and pSub (GFP, green) injection site density heatmap (Figure 1C_1_). (A_2_) a diagonal lat-mMN-to-med-mMN axis of mMN expression (Figure 1B_2_) overlaid with density heatmap of projections from dSub (magenta) and pSub (GFP; Figure 1C_2_). The correspondence between axon intensity at the injection and projection sites and Ten3/Lphn2 expression supports the ‘Ten3→Ten3, Lphn2→Lphn2’ connectivity rule. (B) Additional representative image of topographic projections in lat-mMN and med-mMN (B_1_; from dual dSub and pSub injections), show axons do not segregate until reaching the target and then turn (arrows) to fill Ten3^+^ (B_2_) and Lphn2^+^ subfields (B_3_). B_2_ and B_3_ are from dotted rectangle in B_1_, labeled only with mCherry or GFP channels, respectively. Scale bar, 50 μm. (C) Schematic for *ex vivo* injections for developmental tracing of entorhinal cortex→subiculum projection. (D) Representative images of projection of entorhinal cortex axons into the molecular layer of subiculum in wildtype animals at E17.5 (D_1_), P0 (D_2_), P2 (D_3_), and P6 (D_4_). Axons begin to invade subiculum at E17.5 and progressively increase in density through P6. Scale bar, 100 μm. (E) Representative images of MEC axons (magenta) at P0 entering Ten3-high dSub (cyan) and turning before proceeding to Lphn2-high pSub (yellow) at P0. The three images are from the same confocal section showing axons (E_1_), Ten3 protein (E_2_), Lphn2 protein (E_3_), and merge (E_4_) channels. Scale bar, 50 μm. (F) Representative images of subiculum axon tract at P0 along the path to mMN with extensive intermingling of Ten3^+^ (cyan) and Lphn2^+^ (yellow) axons in a *Ten3^HA/HA^;Lphn2^mVenus/mVenus^* mouse. The three images are from the same confocal section showing Ten3 (F_1_), Lphn2 (F_2_), and merge (F_3_) channels. Scale bar, 50 μm. (G) Representative images of subiculum axon tract at P0, entering mMN target where axons experience Ten3^+^ and Lphn2^+^ subfields simultaneously. Ten3 (cyan) and Lphn2 (yellow) axons remain intermingled in the tract up until terminal projections in the target. The three images are from the same confocal section showing Ten3 (G_1_), Lphn2 (G_2_), and merge (G_3_) channels. Scale bar, 25 μm.

**Figure S2.**
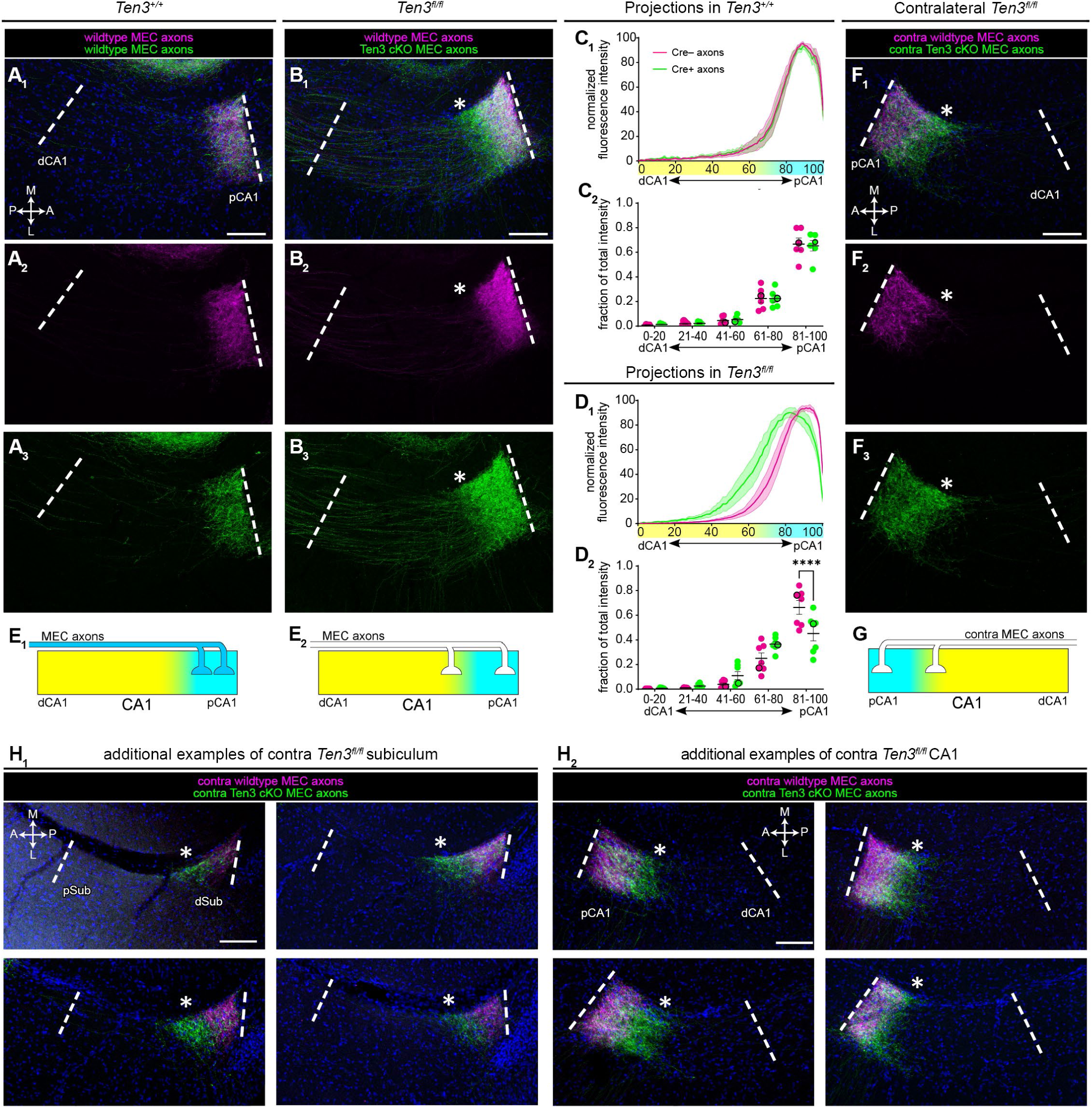
Additional analyses of the MEC axon projection, related to Figure 2. (A) Representative images of projection of Cre^–^ axons (magenta), and Cre^+^ axons (green) into molecular layer of CA1 in *Ten3^+/+^* controls. The three images are from the same confocal section showing merge (A_1_), magenta only (A_2_), or green only (A_3_) channels. Corresponding injection is in Figure 2C. (B) Same as A but for *Ten3^fl/fl^* mice. *Ten3* cKO axons in *Ten3^fl/fl^* mice spread beyond the Cre^–^ control axons towards dCA1 (asterisk, in identical position across three images). Corresponding injection is in Figure 2E. (C_1_) Normalized fluorescence intensity traces of the Cre^–^ (magenta) and Cre^+^ (green) projections along the distal→proximal axis of CA1 in *Ten3^+/+^*controls (n = 6 mice). Injection positions are in Figure 2B. Ten3 (cyan) and Lphn2 (yellow) expression is represented along the x-axis. Mean (dark line) ± SEM (shaded area). (C_2_) Fraction of total projection intensity (same as C_1_) in 20% bins of CA1 axis of the Cre^–^ (magenta) and Cre^+^ (green) projections along the distal–proximal axis of CA1 in *Ten3^+/+^* controls (n = 6 mice). Black outlines indicate representative animal in A. Mean ± SEM. Two-way ANOVA corrected for multiple comparisons (Šídák correction), no significant differences. (D) Same as C but for *Ten3^fl/fl^* (n = 7) mice. **** p < 0.0001. (E) Schematic summary of target selection of MEC axons in CA1 in control (E_1_) and *Ten3^MEC^*-cKO (E_2_) based on results in A–D. (F) Same as B, but for projections into contralateral CA1. Contralateral Cre^+^, *Ten3^MEC^*-cKO axons spread beyond the contralateral Cre^–^ control axons towards dCA1 (asterisk, in identical position across three images). Corresponding injection is in Figure 2K. (G) Schematic summary of target selection of MEC axons in contralateral CA1 in *Ten3^MEC^*-cKO mice. (H) Four additional examples of contralateral projections in subiculum (H_1_) and CA1 (H_2_) of *Ten3^MEC^*-cKO. Asterisks indicate GFP^+^ axon mistargeting. Scale bars, 100 μm.

**Figure S3.**
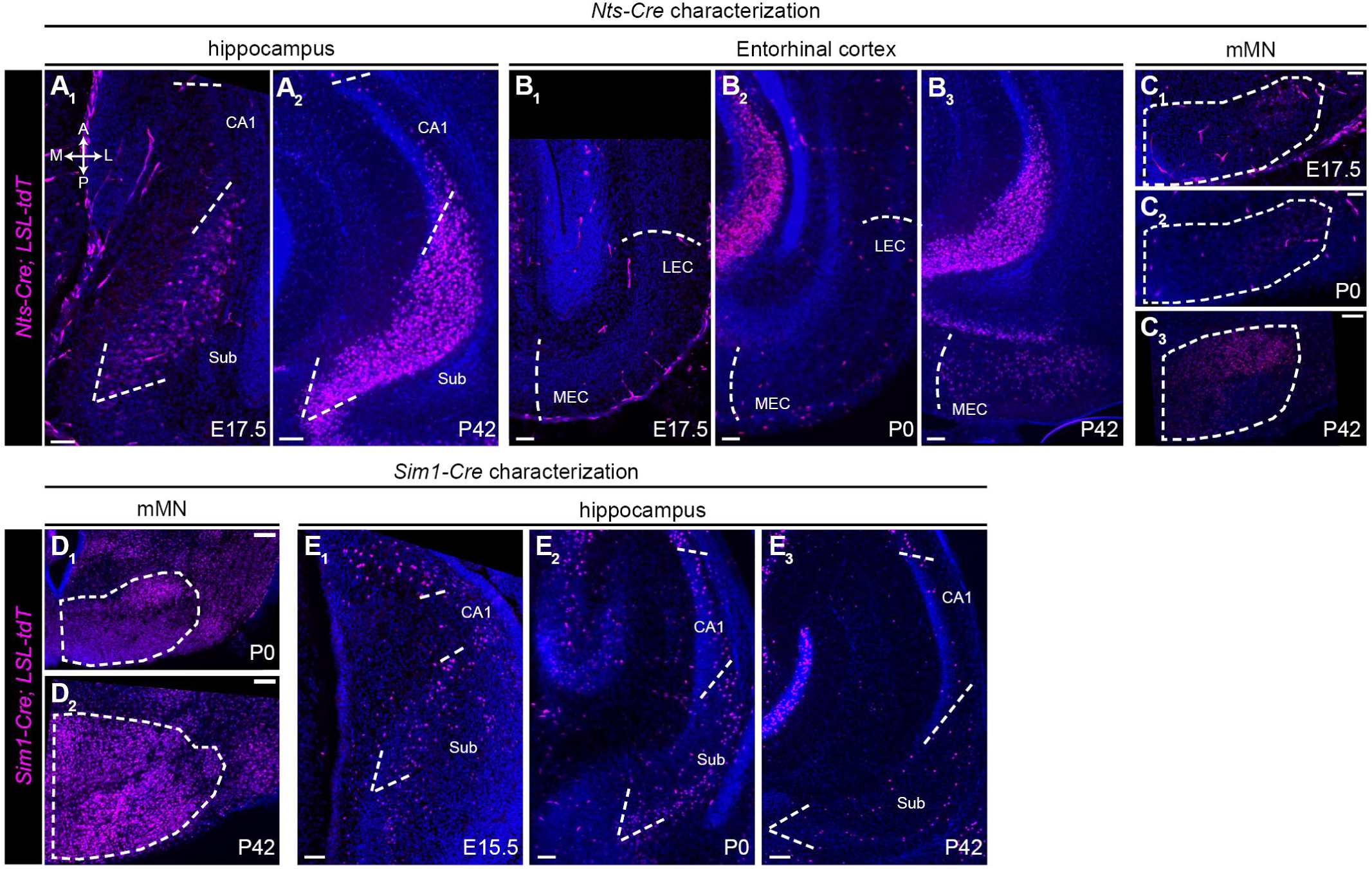
*Nts-Cre* and *Sim1-Cre* characterizations, related to Figures 3–7. (A) Expression of *Nts-Cre* in subiculum and CA1, assessed by a nuclear tdTomato Cre reporter at E17.5 (A_1_) and P42 (A_2_). P0 data is in Figure 3A. The Cre reporter is sparsely expressed in subiculum at E17.5 and is denser by P0. Little CA1 expression is found even in adult. Scale bar, 50 μm (A_1_), 100 μm (A_2_). (B) Same as A, except in entorhinal cortex with the addition of P0 (B_2_). Expression is sparse in entorhinal cortex during development, ensuring target-only deletions in our examination of the MEC→dSub projection (Figure 3). Scale bar, 50 μm (B_1_, B_2_), 100 μm (B_3_). (C) Same as A, except in medial mammillary nucleus (mMN) with the addition of P0 (C_2_). Expression is sparse in mMN during development, ensuring axon-only deletions in our examination of the subiculum→mMN projections (Figures 4 and 6). Scale bar, 50 μm (C_1_, C_2_), 100 μm (C_3_). (D) Expression of *Sim1-Cre* in mMN, assessed by a nuclear tdTomato Cre reporter at P0 (D_1_) and P42 (D_2_). E15.5 data is in Figure 5A. The Cre reporter is densely expressed in mMN by E15.5. Scale bars, 50 μm (D_1_), 100 μm (D_2_). (E) Same as D, except in subiculum with the addition of E15.5 (E_1_). Expression is sparse in subiculum during development and even in adults, ensuring target-only deletions in our examination of the subiculum→mMN projections (Figures 5 and 7). Scale bar, 50 μm (E_1_, E_2_), 100 μm (E_3_). All panels show endogenous tdTomato fluorescence, except for E17.5 *Nts-Cre* panels which are stained with RFP antibody (A_1_, B_1_, C_1_).

**Figure S4.**
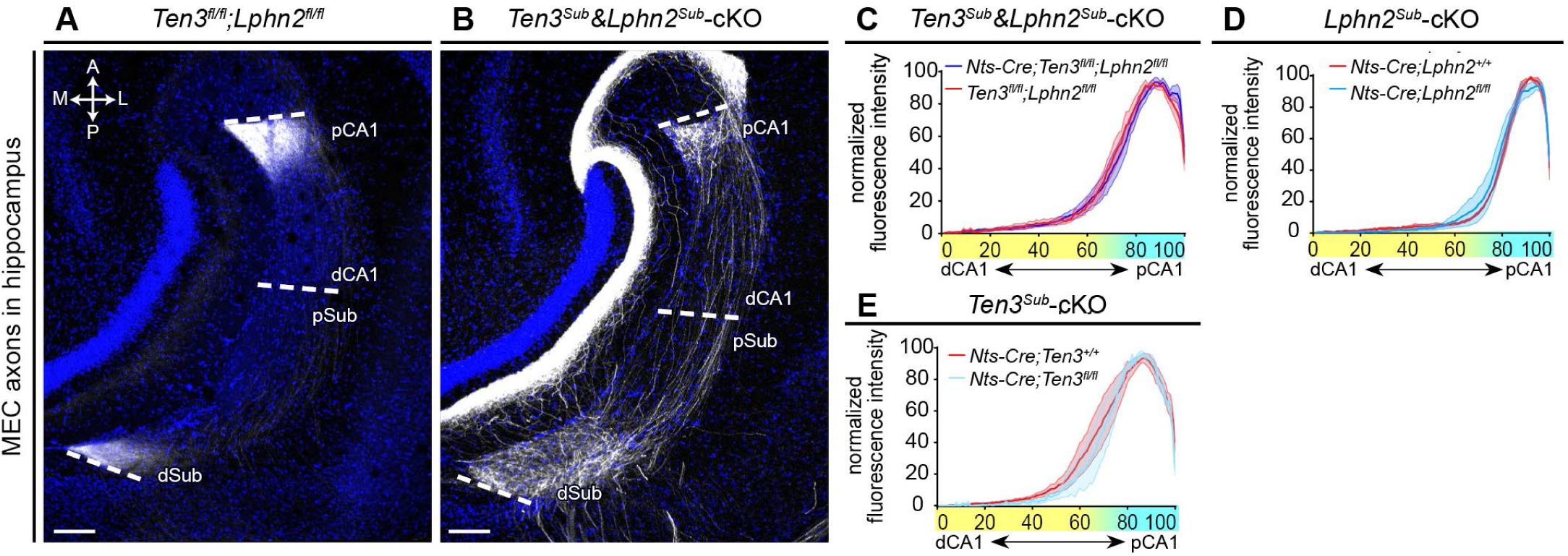
Normal targeting of MEC→pCA1 when *Ten3&Lphn2* were conditionally knocked out in subiculum, related to Figure 3. (A) Representative image of MEC projections into molecular layer of CA1 and subiculum for controls. This is a larger field of view of Figure 3E to show that the same MEC axon population target both subiculum and CA1 with their terminals restricted to dSub and pCA1. Corresponding injection is in Figure 3D. Scale bar, 100 μm. (B) Same as A, but for *Ten3^Sub^&Lphn2^Sub^-*cKO mice. This is a larger field of view of Figure 3G_1_. MEC axons spread towards pSub in subiculum but are restricted to pCA1 in CA1 as in control animals. Corresponding injection is in Figure 3F. (C) Normalized fluorescence intensity traces of MEC axon projections along the distal-to-proximal axis of CA1 internal controls (n = 6; red) and *Ten3^Sub^&Lphn2^Sub^-*cKO mice (n = 5; blue). Injection positions are in Figure 3C. Ten3 (cyan) and Lphn2 (yellow) expression is represented along the x-axis. Mean (dark line) ± SEM (shaded area). (D) Same as C, but for control (n = 3; red) and *Lphn2^Sub^-*cKO (n = 5; blue) mice. (E) Same as C, but for control (n = 5; red) and *Ten3^Sub^-*cKO (n = 4; blue) mice.

**Figure S5.**
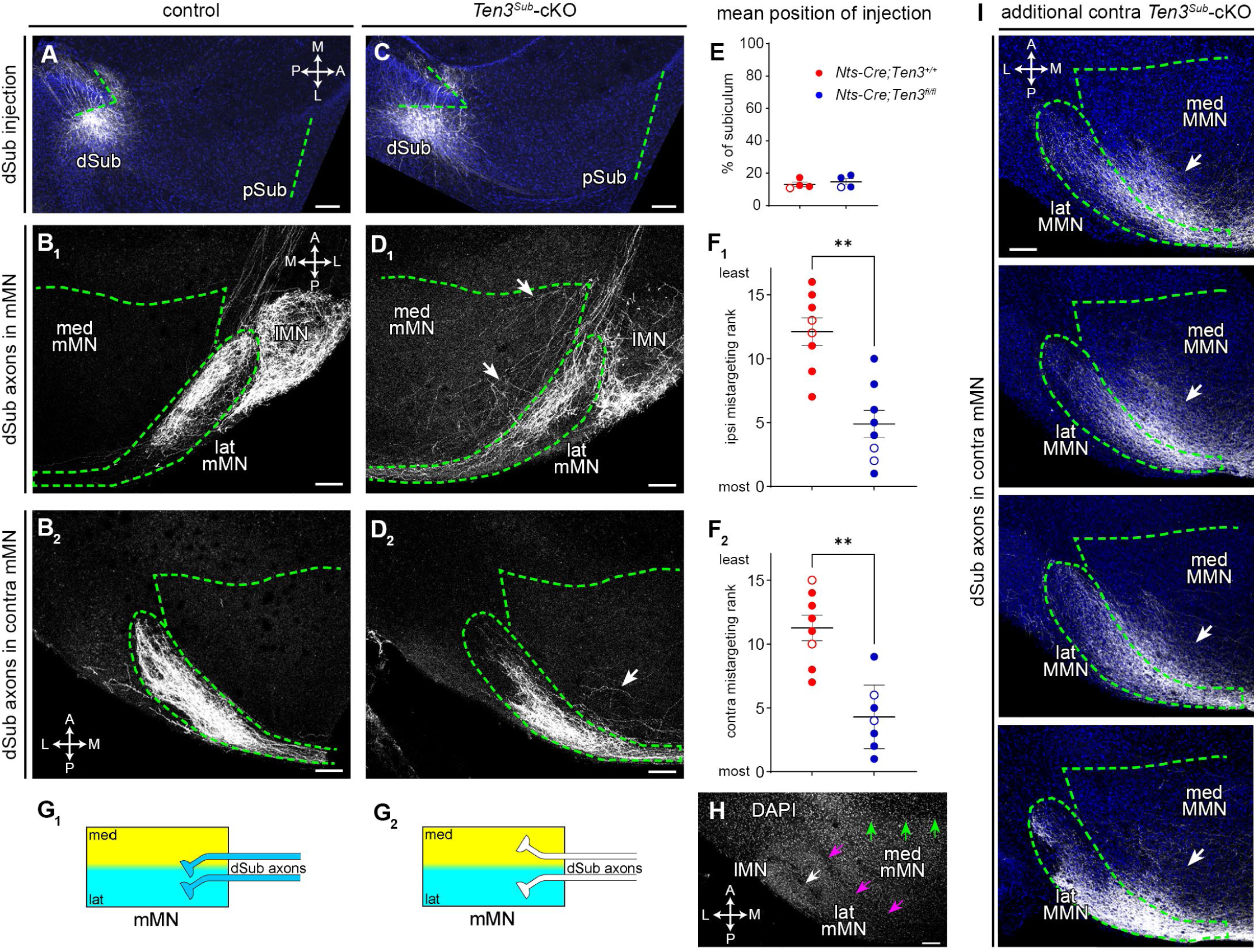
Additional *Ten3* axon deletion data to examine the dSub→lat-mMN projection, related to Figure 4. In an independent set of experiments, we examined the phenotypes of dSub→lat-mMN projection in *Ten3^Sub^*-cKO mice. Here, mistargeting of dSub axons into med-mMN in control and *Ten3^Sub^*-cKO mice were rank-ordered by an experimenter blind to the genotype, based on the relative intensity of dSub axons in med-mMN and lat-mMN (to account for variations in injection intensity of anterograde tracers). (A) Representative image of the *AAV-CreON-mCh* (gray) injection site of controls show the virus is restricted to the most distal region of subiculum. Injection corresponds to animal in (B). (B) Representative images of projection into ipsilateral (B_1_) and contralateral (B_2_) mMN of controls. Labeling in lateral mammillary nucleus (lMN) are unrelated axons from pre-subiculum. (C, D) Same as A, B, but for *Ten3^Sub^*-cKO mice. Even the most distal subiculum axons show aberrant branches in med-mMN (arrows in D). (E) Mean positions of injection sites along the distal-to-proximal axis of subiculum show no differences between control (n = 4, red) and *Ten3^Sub^-*cKO (n = 4, blue). Open circles indicate representative image in A, C. Mean ± SEM. Mann-Whitney test, no significant differences. (F) Rank according to severity of mistargeting for control (n = 8 section, red) and *Ten3^Sub^-*cKO (n = 8 sections, blue) in ipsilateral (F_1_) and contralateral (F_2_) mMN. Contralateral cKO has only n = 7 cKO sections. Open circles indicate representative animals in B, D. Ranked from most severe (1) to least severe (16). Mean ± SEM. Mann-Whitney test, ** p < 0.01. (G) Schematic summary of target selection of dSub axons in contralateral mMN of control (G_1_) and *Ten3^Sub^*-cKO (G_2_) mice based on results in A–F. (H) Representative image of DAPI counterstain in mMN illustrates borders of sub-nuclei used to select regions of interest. Border of lMN (not studied) and lat-mMN (white arrow). Border of Ten3^+^ lat-mMN and Lphn2^+^ med-mMN (magenta arrows). Border of med-mMN and pre-mammillary nucleus (green arrows). Same section as Figure 4I. (I) Four additional examples show mistargeting (arrows) of contralateral dSub→lat-mMN in *Ten3^Sub^*-cKO. Corresponding injections in Figure 4B. Scale bars, 100 μm.

**Figure S6.**
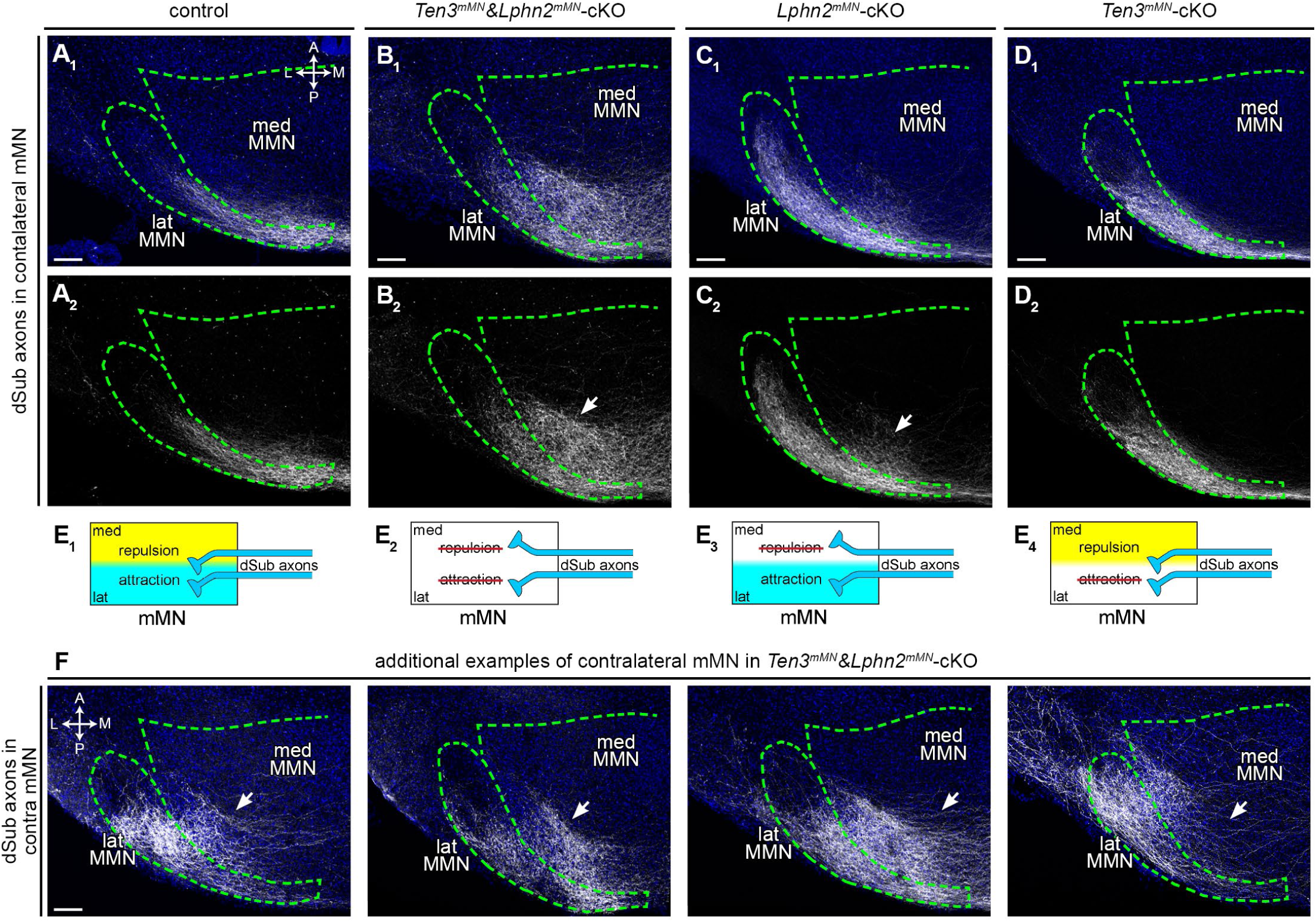
Analysis of contralateral dSub→lat-mMN projections in target deletion conditions, related to Figure 5. (A) Representative images of projection of dSub axons into contralateral medial mammillary nucleus of control mice (A_1_). Bottom panel shows axons without DAPI counterstain (A_2_). Same animal as Figure 5D, E. (B–D) Same as A, but for genotypes indicated above. dSub axons in *Ten3^mMN^&Lphn2^mMN^*-cKO (B) and *Lphn2^mMN^-*cKO (C) mice spread medially outside of contralateral lat-mMN (arrows in B_2_, C_2_). Same as animal of corresponding genotype in Figure 5F–K. (E) Schematic summary of target selection of dSub axons in contralateral mMN in control (E_1_) and when *Ten3*(E_4_), *Lphn2*(E_3_), or both (E_2_) were conditionally knocked out in mMN. (F) Four additional examples show mistargeting (arrows) of contralateral dSub→lat-mMN projections in *Ten3^mMN^&Lphn2^mMN^*-cKO. Corresponding injections in Figure 5C. Scale bars, 100 μm.

**Figure S7.**
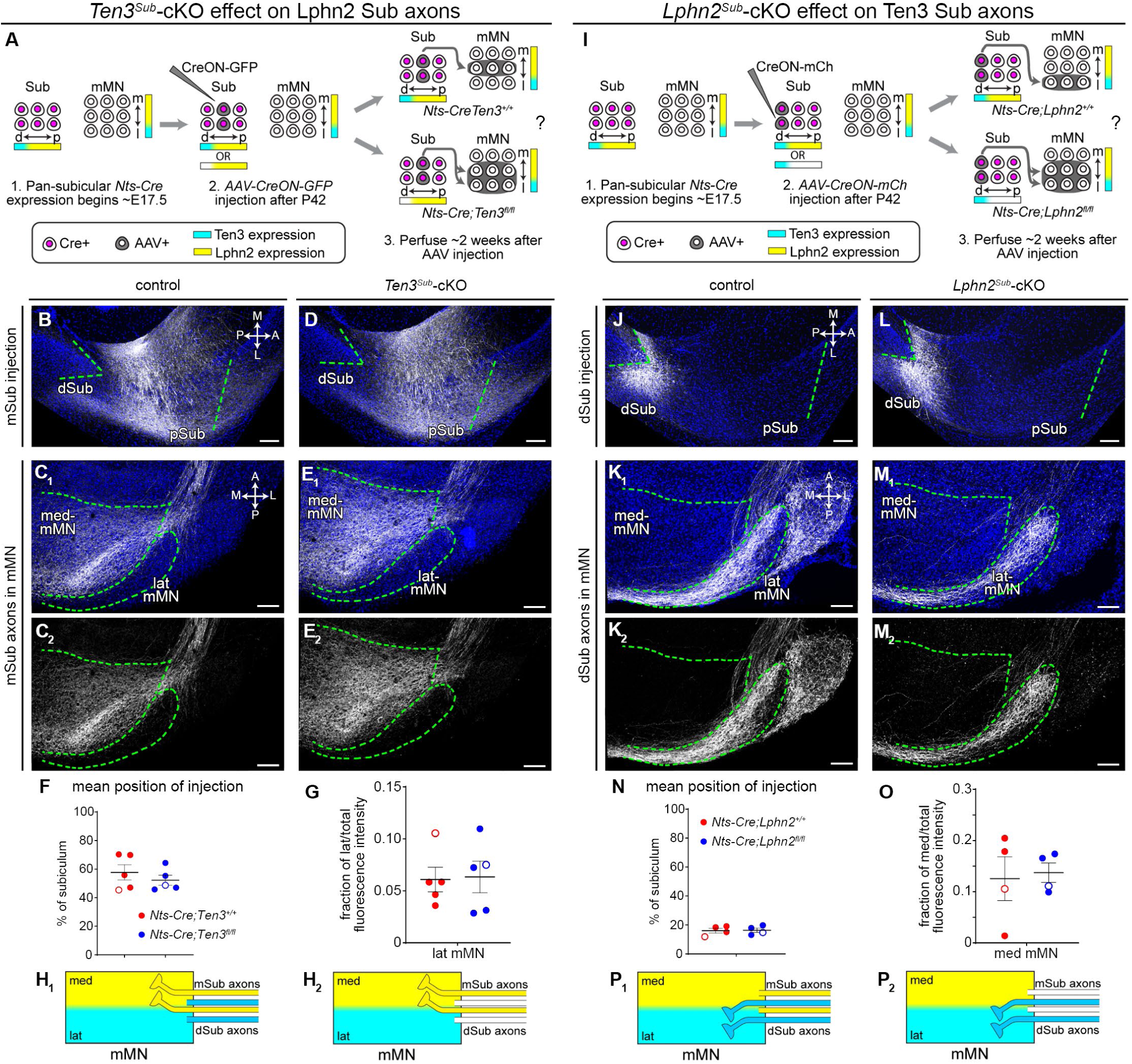
Lack of evidence for axon-axon interaction between Ten3-high dSub and Lphn2-high pSub axons in target selection of mMN, related to Figures 4 and 6. (A) Injection strategy and potential results for tracing functionally wildtype pSub (Lphn2-high) axons into medial mammillary nucleus (mMN) of *Nts-Cre;Ten3^+/+^* control (top right) and *Ten3^Sub^*-cKO mice (bottom right). (B) Representative image of the *AAV-CreON-GFP* (gray) injection site of controls show the virus is restricted to proximal region of subiculum. Injection corresponds to animal in (C). (C) Representative images of projection of pSub axons (gray) into mMN of controls (C_1_). Bottom panel shows axons without DAPI counterstain (C_2_). (D, E) Same as B, C, but for *Ten3^Sub^*-cKO. Lphn2-high pSub axon targeting is unaffected by *Ten3* deletion in subiculum axons. (F) Mean positions of injection sites along the proximal-distal axis of subiculum show no differences between controls (n = 5, red) and *Ten3^Sub^*-cKO (n = 5, blue). Open circles indicate representative animals in B, D. Mean ± SEM. Mann-Whitney. (G) Fraction of total projection intensity in lat-mMN of the pSub axons in controls (n = 5, red) and *Ten3^Sub^*-cKO (n = 5, blue). Open circles indicate representative animals in C, E. Mean ± SEM. Mann-Whitney, no significant differences. (H) Schematic summary of target selection of pSub axons in mMN of control (H_1_) and *Ten3^Sub^*-cKO (H_2_) mice based on results in B–G. For clarity, we truncated the dSub axons in which *Ten3* was deleted. (I–P) Same as A–H, but for functionally wildtype dSub (Ten3-high) *AAV-CreON-mCh* (gray) injections into *Nts-Cre;Lphn2^+/+^* control (n = 4, red) and *Lphn2^Sub^*-cKO mice (n = 4, blue). Mean ± SEM. Mann-Whitney, no significant differences. Scale bars, 100 μm.

**Table S1.**
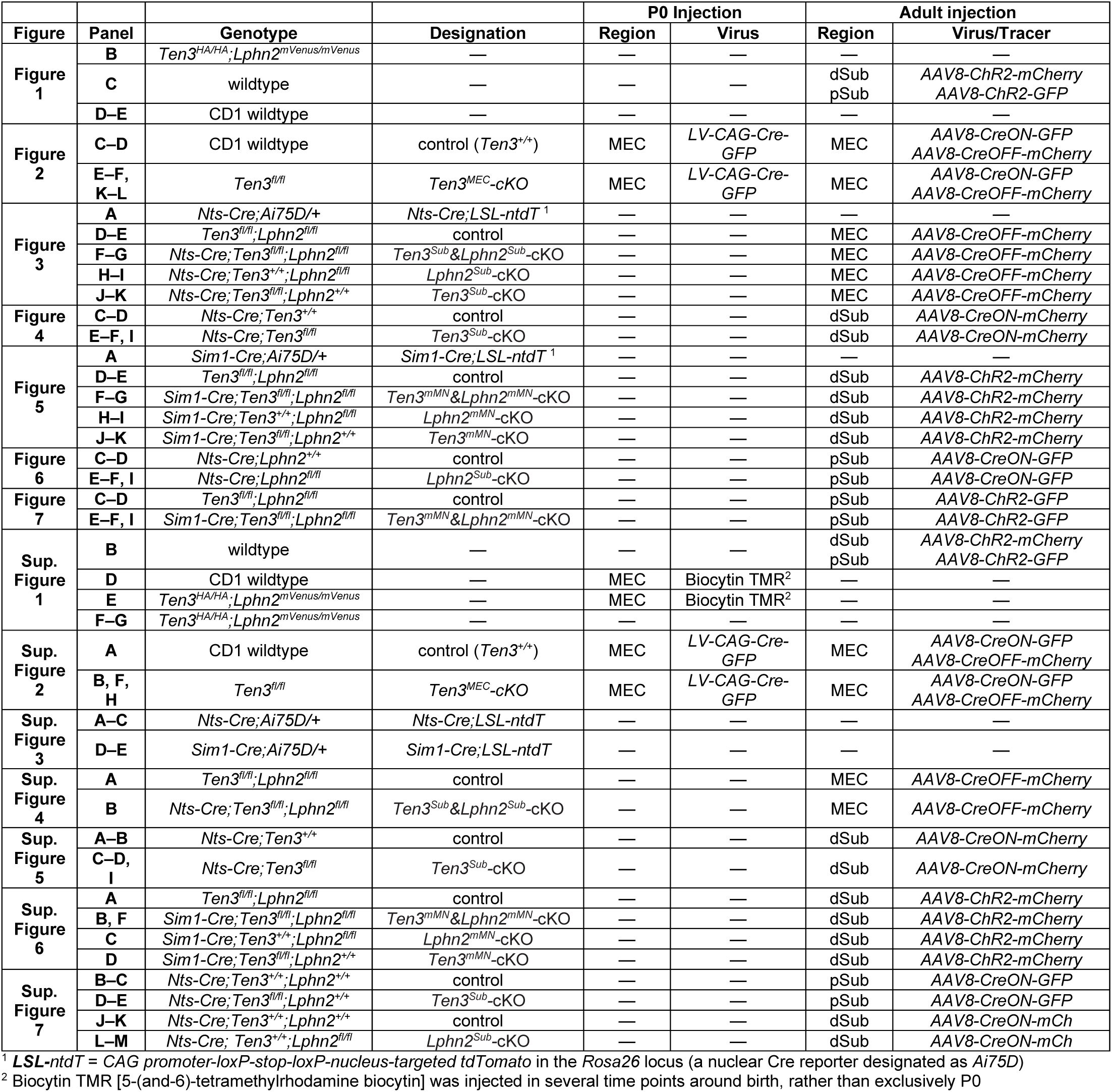
Summary of genotypes, viruses, and injections in each experiment, arranged according to figure panel, related to Figures 1–7 and Figures S1–S7.

## Notes

### Competing Interest Statement

The authors have declared no competing interest.

### Summary of Updates

Additional discussion and supplemental data was added.

## REFERENCES

1. Sperry, R.W. (1963). Chemoaffinity in the orderly growth of nerve fiber patterns and connections*. Proc. Natl. Acad. Sci. 50, 703–710. 10.1073/pnas.50.4.703.

2. Sanes, J.R., and Zipursky, S.L. (2020). Synaptic Specificity, Recognition Molecules, and Assembly of Neural Circuits. Cell 181, 536–556. 10.1016/j.cell.2020.04.008.

3. Kolodkin, A.L., and Tessier-Lavigne, M. (2011). Mechanisms and Molecules of Neuronal Wiring: A Primer. Cold Spring Harb. Perspect. Biol. 3, a001727. 10.1101/cshperspect.a001727.

4. Walter, J., Kern-Veits, B., Huf, J., Stolze, B., and Bonhoeffer, F. (1987). Recognition of position-specific properties of tectal cell membranes by retinal axons in vitro. Development 101, 685–696.

5. Cheng, H.-J., Nakamoto, M., Bergemann, A.D., and Flanagan, J.G. (1995). Complementary gradients in expression and binding of ELF-1 and Mek4 in development of the topographic retinotectal projection map. Cell 82, 371–381. 10.1016/0092-8674(95)90426-3.

6. Drescher, U., Kremoser, C., Handwerker, C., Löschinger, J., Noda, M., and Bonhoeffer, F. (1995). In vitro guidance of retinal ganglion cell axons by RAGS, a 25 kDa tectal protein related to ligands for Eph receptor tyrosine kinases. Cell 82, 359–370. 10.1016/0092-8674(95)90425-5.

7. Hong, W., Mosca, T.J., and Luo, L. (2012). Teneurins instruct synaptic partner matching in an olfactory map. Nature 484, 201–207. 10.1038/nature10926.

8. Berns, D.S., DeNardo, L.A., Pederick, D.T., and Luo, L. (2018). Teneurin-3 controls topographic circuit assembly in the hippocampus. Nature 554, 328–333. 10.1038/nature25463.

9. Pederick, D.T., Lui, J.H., Gingrich, E.C., Xu, C., Wagner, M.J., Liu, Y., He, Z., Quake, S.R., and Luo, L. (2021). Reciprocal repulsions instruct the precise assembly of parallel hippocampal networks. Science 372, 1068–1073. 10.1126/science.abg1774.

10. Duan, X., Krishnaswamy, A., Laboulaye, M.A., Liu, J., Peng, Y.-R., Yamagata, M., Toma, K., and Sanes, J.R. (2018). Cadherin Combinations Recruit Dendrites of Distinct Retinal Neurons to a Shared Interneuronal Scaffold. Neuron 99, 1145–1154.e6. 10.1016/j.neuron.2018.08.019.

11. Dombrovski, M., Zang, Y., Frighetto, G., Vaccari, A., Jang, H., Mirshahidi, P.S., Xie, F., Sanfilippo, P., Hina, B.W., Rehan, A., et al. (2025). Molecular gradients shape synaptic specificity of a visuomotor transformation. Nature 644, 453–462. 10.1038/s41586-025-09037-4.

12. Li, Z., Lyu, C., Xu, C., Hu, Y., Luginbuhl, D.J., Caspi-Lebovic, A.B., Priest, J.M., Özkan, E., and Luo, L. (2025). Repulsive interactions instruct synaptic partner matching in an olfactory circuit. Preprint at bioRxiv, 10.1101/2025.03.01.640985 https://doi.org/10.1101/2025.03.01.640985.

13. Lyu, C., Li, Z., Xu, C., Kalai, J., and Luo, L. (2026). Rewiring an olfactory circuit by altering the combinatorial code of cell-surface proteins. Preprint at bioRxiv, 10.1101/2025.03.01.640986 https://doi.org/10.1101/2025.03.01.640986.

14. Azevedo, F.A.C., Carvalho, L.R.B., Grinberg, L.T., Farfel, J.M., Ferretti, R.E.L., Leite, R.E.P., Jacob Filho, W., Lent, R., and Herculano-Houzel, S. (2009). Equal numbers of neuronal and nonneuronal cells make the human brain an isometrically scaled-up primate brain. J. Comp. Neurol. 513, 532–541. 10.1002/cne.21974.

15. Andrade-Moraes, C.H., Oliveira-Pinto, A.V., Castro-Fonseca, E., da Silva, C.G., Guimarães, D.M., Szczupak, D., Parente-Bruno, D.R., Carvalho, L.R.B., Polichiso, L., Gomes, B.V., et al. (2013). Cell number changes in Alzheimer’s disease relate to dementia, not to plaques and tangles. Brain J. Neurol. 136, 3738–3752. 10.1093/brain/awt273.

16. Goriely, A. (2025). Eighty-six billion and counting: do we know the number of neurons in the human brain? Brain 148, 689–691. 10.1093/brain/awae390.

17. Bausch-Fluck, D., Goldmann, U., Müller, S., van Oostrum, M., Müller, M., Schubert, O.T., and Wollscheid, B. (2018). The in silico human surfaceome. Proc. Natl. Acad. Sci. 115, E10988–E10997. 10.1073/pnas.1808790115.

18. Luo, L. (2021). Principles of Neurobiology 2nd ed. (CRC Press).

19. Cang, J., and Feldheim, D.A. (2013). Developmental Mechanisms of Topographic Map Formation and Alignment. Annu. Rev. Neurosci. 36, 51–77. 10.1146/annurev-neuro-062012-170341.

20. Brown, A., Yates, P.A., Burrola, P., Ortuño, D., Vaidya, A., Jessell, T.M., Pfaff, S.L., O’Leary, D.D.M., and Lemke, G. (2000). Topographic Mapping from the Retina to the Midbrain Is Controlled by Relative but Not Absolute Levels of EphA Receptor Signaling. Cell 102, 77–88. 10.1016/S0092-8674(00)00012-X.

21. Feldheim, D.A., Kim, Y.-I., Bergemann, A.D., Frisén, J., Barbacid, M., and Flanagan, J.G. (2000). Genetic Analysis of Ephrin-A2 and Ephrin-A5 Shows Their Requirement in Multiple Aspects of Retinocollicular Mapping. Neuron 25, 563–574. 10.1016/S0896-6273(00)81060-0.

22. Rashid, T., Upton, A.L., Blentic, A., Ciossek, T., Knöll, B., Thompson, I.D., and Drescher, U. (2005). Opposing Gradients of Ephrin-As and EphA7 in the Superior Colliculus Are Essential for Topographic Mapping in the Mammalian Visual System. Neuron 47, 57–69. 10.1016/j.neuron.2005.05.030.

23. Feldheim, D.A., Vanderhaeghen, P., Hansen, M.J., Frisén, J., Lu, Q., Barbacid, M., and Flanagan, J.G. (1998). Topographic Guidance Labels in a Sensory Projection to the Forebrain. Neuron 21, 1303–1313. 10.1016/S0896-6273(00)80650-9.

24. Cang, J., Kaneko, M., Yamada, J., Woods, G., Stryker, M.P., and Feldheim, D.A. (2005). Ephrin-As Guide the Formation of Functional Maps in the Visual Cortex. Neuron 48, 577–589. 10.1016/j.neuron.2005.10.026.

25. Pfeiffenberger, C., Yamada, J., and Feldheim, D.A. (2006). Ephrin-As and Patterned Retinal Activity Act Together in the Development of Topographic Maps in the Primary Visual System. J. Neurosci. 26, 12873–12884. 10.1523/JNEUROSCI.3595-06.2006.

26. Suetterlin, P., and Drescher, U. (2014). Target-Independent EphrinA/EphA-Mediated Axon-Axon Repulsion as a Novel Element in Retinocollicular Mapping. Neuron 84, 740–752. 10.1016/j.neuron.2014.09.023.

27. Igarashi, K.M., Ito, H.T., Moser, E.I., and Moser, M.-B. (2014). Functional diversity along the transverse axis of hippocampal area CA1. FEBS Lett. 588, 2470–2476. 10.1016/j.febslet.2014.06.004.

28. Cembrowski, M.S., Phillips, M.G., DiLisio, S.F., Shields, B.C., Winnubst, J., Chandrashekar, J., Bas, E., and Spruston, N. (2018). Dissociable Structural and Functional Hippocampal Outputs via Distinct Subiculum Cell Classes. Cell 173, 1280–1292.e18. 10.1016/j.cell.2018.03.031.

29. Tamamaki, N., and Nojyo, Y. (1995). Preservation of topography in the connections between the subiculum, field CA1, and the entorhinal cortex in rats. J. Comp. Neurol. 353, 379–390. 10.1002/cne.903530306.

30. Allen, G.V., and Hopkins, D.A. (1989). Mamillary body in the rat: topography and synaptology of projections from the subicular complex, prefrontal cortex, and midbrain tegmentum. J. Comp. Neurol. 286, 311–336. 10.1002/cne.902860303.

31. Shibata, H. (1989). Descending projections to the mammillary nuclei in the rat, as studied by retrograde and anterograde transport of wheat germ agglutinin-horseradish peroxidase. J. Comp. Neurol. 285, 436–452. 10.1002/cne.902850403.

32. Kishi, T., Tsumori, T., Ono, K., Yokota, S., Ishino, H., and Yasui, Y. (2000). Topographical organization of projections from the subiculum to the hypothalamus in the rat. J. Comp. Neurol. 419, 205–222. https://doi.org/10.1002/(SICI)1096-9861(20000403)419:2%253C205::AID-CNE5%253E3.0.CO;2-0.

33. Naber, P.A., Lopes da Silva, F.H., and Witter, M.P. (2001). Reciprocal connections between the entorhinal cortex and hippocampal fields CA1 and the subiculum are in register with the projections from CA1 to the subiculum. Hippocampus 11, 99–104. 10.1002/hipo.1028.

34. van Strien, N.M., Cappaert, N.L.M., and Witter, M.P. (2009). The anatomy of memory: an interactive overview of the parahippocampal–hippocampal network. Nat. Rev. Neurosci. 10, 272–282. 10.1038/nrn2614.

35. Wright, N.F., Erichsen, J.T., Vann, S.D., O’Mara, S.M., and Aggleton, J.P. (2010). Parallel but separate inputs from limbic cortices to the mammillary bodies and anterior thalamic nuclei in the rat. J. Comp. Neurol. 518, 2334–2354. 10.1002/cne.22336.

36. Pederick, D.T., Perry-Hauser, N.A., Meng, H., He, Z., Javitch, J.A., and Luo, L. (2023). Context-dependent requirement of G protein coupling for Latrophilin-2 in target selection of hippocampal axons. eLife 12, e83529. 10.7554/eLife.83529.

37. Donohue, J.D., Amidon, R.F., Murphy, T.R., Wong, A.J., Liu, E.D., Saab, L., King, A.J., Pae, H., Ajayi, M.T., and Anderson, G.R. (2021). Parahippocampal latrophilin-2 (ADGRL2) expression controls topographical presubiculum to entorhinal cortex circuit connectivity. Cell Rep. 37, 110031. 10.1016/j.celrep.2021.110031.

38. Liakath-Ali, K., Refaee, R., and Südhof, T.C. (2024). Cartography of teneurin and latrophilin expression reveals spatiotemporal axis heterogeneity in the mouse hippocampus during development. PLOS Biol. 22, e3002599. 10.1371/journal.pbio.3002599.

39. Chon, U., Pederick, D.T., Song, J.H., Zhang, Y., Rana, I., and Luo, L. (2025). Inverse expression of Ten3 and Lphn2 across the developing mouse brain reveals a global strategy for circuit assembly. Preprint at bioRxiv, 10.1101/2025.08.13.670004 https://doi.org/10.1101/2025.08.13.670004.

40. Anderson, G.R., Maxeiner, S., Sando, R., Tsetsenis, T., Malenka, R.C., and Südhof, T.C. (2017). Postsynaptic adhesion GPCR latrophilin-2 mediates target recognition in entorhinal-hippocampal synapse assembly. J. Cell Biol. 216, 3831–3846. 10.1083/jcb.201703042.

41. Wang, F., Flanagan, J., Su, N., Wang, L.-C., Bui, S., Nielson, A., Wu, X., Vo, H.-T., Ma, X.-J., and Luo, Y. (2012). RNAscope: A Novel in Situ RNA Analysis Platform for Formalin-Fixed, Paraffin-Embedded Tissues. J. Mol. Diagn. 14, 22–29. 10.1016/j.jmoldx.2011.08.002.

42. The MouseLight Project (Howard Hughes Medical Institute, Janelia Campus).

43. Digital Brain - Projectome Atlas (Institute of Neuroscience, Chinese Academy of Sciences).

44. Qiu, S., Hu, Y., Huang, Y., Gao, T., Wang, X., Wang, D., Ren, B., Shi, X., Chen, Y., Wang, X., et al. (2024). Whole-brain spatial organization of hippocampal single-neuron projectomes. Science 383, eadj9198. 10.1126/science.adj9198.

45. Winnubst, J., Bas, E., Ferreira, T.A., Wu, Z., Economo, M.N., Edson, P., Arthur, B.J., Bruns, C., Rokicki, K., Schauder, D., et al. (2019). Reconstruction of 1,000 Projection Neurons Reveals New Cell Types and Organization of Long-Range Connectivity in the Mouse Brain. Cell 179, 268–281.e13. 10.1016/j.cell.2019.07.042.

46. Economo, M.N., Clack, N.G., Lavis, L.D., Gerfen, C.R., Svoboda, K., Myers, E.W., and Chandrashekar, J. (2016). A platform for brain-wide imaging and reconstruction of individual neurons. eLife 5, e10566. 10.7554/eLife.10566.

47. Boucard, A.A., Maxeiner, S., and Südhof, T.C. (2014). Latrophilins Function as Heterophilic Cell-adhesion Molecules by Binding to Teneurins: REGULATION BY ALTERNATIVE SPLICING*. J. Biol. Chem. 289, 387–402. 10.1074/jbc.M113.504779.

48. Imai, T., Yamazaki, T., Kobayakawa, R., Kobayakawa, K., Abe, T., Suzuki, M., and Sakano, H. (2009). Pre-target axon sorting establishes the neural map topography. Science 325, 585–590. 10.1126/science.1173596.

49. Sando, R., Jiang, X., and Südhof, T.C. (2019). Latrophilin GPCRs direct synapse specificity by coincident binding of FLRTs and teneurins. Science 363, eaav7969. 10.1126/science.aav7969.

50. Sando, R., and Südhof, T.C. (2021). Latrophilin GPCR signaling mediates synapse formation. eLife 10, e65717. 10.7554/eLife.65717.

51. Sangster, K.T., Zhang, X., Toro, D. del, Sarantopoulos, C., Moses, A.M., Mahasenan, S., Pederick, D.T., Roome, R.B., Seiradake, E., Luo, L., et al. (2025). Teneurin-3 and Latrophilin-2 are required for the formation of a somatotopic map and somatosensory topognosis. Preprint at bioRxiv, 10.1101/2025.08.13.670179 https://doi.org/10.1101/2025.08.13.670179.

52. Luo, L. (2021). Architectures of neuronal circuits. Science 373, eabg7285. 10.1126/science.abg7285.

53. Sun, Y., Pederick, D.T., Xu, X., Luo, L., and Giocomo, L.M. (2026). Topographic CA1 input shapes subicular spatial coding. Preprint at bioRxiv, 10.64898/2026.03.24.714092 https://doi.org/10.64898/2026.03.24.714092.

54. Egea, J., and Klein, R. (2007). Bidirectional Eph–ephrin signaling during axon guidance. Trends Cell Biol. 17, 230–238. 10.1016/j.tcb.2007.03.004.

55. Fenno, L.E., Mattis, J., Ramakrishnan, C., Hyun, M., Lee, S.Y., He, M., Tucciarone, J., Selimbeyoglu, A., Berndt, A., Grosenick, L., et al. (2014). Targeting cells with single vectors using multiple-feature Boolean logic. Nat. Methods 11, 763–772. 10.1038/nmeth.2996.

56. Saunders, A., Johnson, C., and Sabatini, B. (2012). Novel recombinant adeno-associated viruses for Cre activated and inactivated transgene expression in neurons. Front. Neural Circuits 6. 10.3389/fncir.2012.00047.

57. Boyden, E.S., Zhang, F., Bamberg, E., Nagel, G., and Deisseroth, K. (2005). Millisecond-timescale, genetically targeted optical control of neural activity. Nat. Neurosci. 8, 1263–1268. 10.1038/nn1525.

58. Leinninger, G.M., Opland, D.M., Jo, Y.-H., Faouzi, M., Christensen, L., Cappellucci, L.A., Rhodes, C.J., Gnegy, M.E., Becker, J.B., Pothos, E.N., et al. (2011). Leptin Action via Neurotensin Neurons Controls Orexin, the Mesolimbic Dopamine System and Energy Balance. Cell Metab. 14, 313–323. 10.1016/j.cmet.2011.06.016.

59. Balthasar, N., Dalgaard, L.T., Lee, C.E., Yu, J., Funahashi, H., Williams, T., Ferreira, M., Tang, V., McGovern, R.A., Kenny, C.D., et al. (2005). Divergence of Melanocortin Pathways in the Control of Food Intake and Energy Expenditure. Cell 123, 493–505. 10.1016/j.cell.2005.08.035.

60. Quina, L.A., Harris, J., Zeng, H., and Turner, E.E. (2017). Specific connections of the interpeduncular subnuclei reveal distinct components of the habenulopeduncular pathway. J. Comp. Neurol. 525, 2632–2656. 10.1002/cne.24221.

